# Independent inhibitory control mechanisms for aggressive motivation and action

**DOI:** 10.1101/2022.12.26.521956

**Authors:** Tomohito Minakuchi, Eartha Mae Guthman, Preeta Acharya, Justin Hinson, Weston Fleming, Ilana B. Witten, Stefan N. Oline, Annegret L. Falkner

## Abstract

Social behaviors, like other motivated behaviors, frequently consist of a flexible motivated-seeking or approach phase followed by social action. Dysregulated social behavior may arise from changes to motivation, wherein individuals fail to enter a motivated seeking state, or may be in the execution of the social action itself. However, it is unclear how the brain generates and gates this flexible motivation-to-action sequence, and whether aggressive motivation and action are controlled by separate circuit mechanisms. Here, we record populations of neurons in the ventromedial hypothalamus ventrolateral area (VMHvl) of male mice at cellular resolution during “free” aggression and also during an aggression operant task, where the behaviors that precede attack are stereotyped. We find that this population encodes the temporal sequence of aggressive motivation to action and that the temporal selectivity of neurons is invariant to differences in motivated behavior. To test whether motivation and action could be independently regulated, we focused on two key inhibitory inputs to the VMHvl: a source of local inhibition (VMHvl shell) and the primary source of long-range inhibition (the medial preoptic area, MPO). While we find that the VMHvl receives broad monosynaptic inhibitory input from both inputs, optogenetic perturbation of these inputs during recording reveals temporal selectivity during aggressive motivation and action, suggesting specificity of function. Encoding models applied to population calcium recordings of these inhibitory inputs during naturalistic social interactions and during the social operant task further reveal that these inputs have different temporal dynamics during aggression: VMHvl shell^vgat+^ activity peaks at the start of aggressive interactions, while MPO-VMHvl^vgat+^ activity peaks at behaviorally aligned endpoints of aggressive interactions. Finally, using closed-loop optogenetic stimulation timed to specific phases of the aggression-operant task, we find a double-dissociation of the effects on aggressive motivation and action: activation of MPO-VMHvl^vgat+^, even briefly and temporally distant from the initiation of aggression, produces long-lasting motivational deficits, delaying the initiation of aggression and generating behaviors consistent with an unmotivated state. In contrast, activation of VMHvl shell^vgat+^ produces acute action-related deficits, causing an exit from an attack state. Fitting a Hidden Markov Model (HMM) to behavior further corroborates these findings by showing that MPO-VMHvl^vgat+^ stimulation prolongs a low motivation state and VMHvl shell^vgat+^ promotes exit from an attack state. Together, these data demonstrate how separable inhibitory circuits in the hypothalamus can independently gate the motivational and action phases of aggression through a single locus of control.

## Introduction

Two major questions in neuroscience are first how the brain generates and maintains socially motivated states and second how those states are transformed into action. Examples of common socially motivated states include sexually motivated or aggressively motivated seeking states, in which individuals may seek out opposite sex or same sex conspecifics with the intention of performing specific social actions. Dysregulated social behavior, including aggression, can result from changes to neural circuits that participate in the execution of social actions, or dysregulation can affect neural mechanisms that determine whether highly motivated states will be initiated or reinitiated. Many diverse psychiatric disorders with varied etiologies can result in dysregulated aggression^1,2^, further suggesting that aggression may have multiple types of “brakes” that can go awry. While several brain regions have been previously identified as broadly participating in the gating of aggressive behavior^3–6^, it is currently unclear whether these brakes have separable functions on aggressive motivation and action.

One limitation to previous studies is that aggressive motivation is difficult to quantify. Aggressive motivation can be broadly defined as the internal state that drives animals to seek out opportunities to perform aggressive actions, primarily by approaching potential attack targets. Behaviors that delineate social motivation from action during naturalistic conditions (often referred to as “appetitive” or “preparatory”) are flexible, not easily identified with video, or may be fast and overlapping. For example, during naturalistic social behaviors, the motivated phase may include distinct aggression preparatory behaviors such as approach, investigation and chasing, but these behaviors may not be distinct from each other if animals are in close proximity^7^. In contrast to aggression during free interaction, behavior assays designed to separate the motivational phase from the action phase, including aggression operant tasks, are useful in detecting neurons active during motivation, since the preparatory behavior is stereotyped and easy to quantify^8,9^. An important first step in understanding the neural control of aggressive motivation and action is detecting whether neural circuits that encode motivation-related signals are invariant to these underlying patterns of action, or whether the same neurons represent a more abstract behaviorally invariant motivation-related signal.

The ventromedial hypothalamus ventrolateral area (VMHvl) has emerged in the last decade as a key region in the mammalian neural circuit for aggression. Neurons in this brain area respond both during the sensory investigation of a conspecific and during aggressive action^10,11^, and stimulation of this brain region promotes attack in both sexes^11–13^. In addition to its role in attack, the VMHvl is also involved in promoting an aggressively motivated state: stimulation during the motivated seeking phase during an aggression operant task increases aggression-seeking^8^. However, since the motivational phase of aggression can flexibly unfold with different behaviors and across different timescales, it is unclear whether neurons encode a more abstract motivated internal state, or are yoked to specific motivational behaviors. The “core” of the VMHvl (historically the focus of previous neural recordings during aggression) is a densely glutamatergic with few intermingled inhibitory neurons. Connectivity within the VMHvl core has been recently shown to be “ultrasparse”, with only very rare synaptic coupling between neighboring neurons^14^. This constraint suggests that not only do VMHvl neurons likely inherit their activity profiles from their inputs, but also that inputs to these neurons may target specific subsets of neurons in the VMHvl without their influence spreading to neighboring neurons, allowing them to perform separable functions. Neurons in the VMHvl core receive a wide variety of inputs, with many of the most prominent being long-range inhibitory inputs from hypothalamic and extrahypothalamic structures^15^. The largest of these inputs is from the medial preoptic area (MPO), a complex structure with established roles in many non-aggressive social behaviors including sexual and parental behaviors^16,17^. In addition to these inputs, an understudied source of local inhibition at the ventral and lateral edges of the VMHvl (the VMHvl shell) has been shown to be able to gate aggression by acting through the VMHvl core^18,19^, though little is known about the activity profile of these neurons during aggressive motivation or action. These inputs could potentially exert preferential control of aggressive motivation or action either through specific projections to subsets of neurons in the VMHvl (motivation or action specific labeled lines), or may project broadly and influence motivation and action through the temporal selectivity of their activity patterns, creating motivation or action biased gates.

Here, using a combination of *in vivo* chronic extracellular recording, cell-type-specific calcium recording, *ex vivo* recordings, and optogenetic perturbation, we find evidence for broad connectivity, but temporally selective patterns of activity during aggressive motivation and action. We find that neurons in the VMHvl encode the temporal sequence of behavior of motivation to action, even when the sequence of motivational behaviors is different. In addition, we find inhibitory neurons in the VMHvl shell and inhibitory projections from the MPO have distinct activity profiles during aggressive motivation and actions, and activation of these inputs preferentially gates aggressive motivation and action, respectively. Overall, these data suggest that the VMHvl does not simply represent a unidimensional signal of aggressiveness, but instead encodes independently gated temporal phases of aggression.

## Results

In order to compare the neural dynamics under conditions where the motivational phase of aggression contains different actions, we perform two behavioral tests of aggression in the same session: “free” aggression and an aggression operant task where a stereotyped action (a nosepoke) is required for access to aggression (Supplementary Fig 1a). For tests of free aggression we allow male mice to interact repeatedly with previously-defeated adult males in the homecage (Fig 1a) and interleave these tests with interactions with females and novel objects. To generate stereotyped motivation-related actions, we trained mice on a novel automated version of our previously developed^8^ aggression-operant task (“Social Operant with Actuator-delivered Reward”, SOAR, Fig 1b). Use of this task allows us first to constrain the actions performed during the motivational phase of aggression, and also to stretch this process out in time. During the SOAR task, animals are given access to a 2-port nose-poke panel in their homecage whereby activating the “social” poke gives animals brief access (via an automated linear actuator) to a tethered and previously-defeated male that they can attack for a brief duration (the “dwell” period). The descent of the actuator is triggered by the nosepoke following a random trial-to-trial delay between 0-1s. The epoch of social interaction (“dwell”) times is randomized to be between 1-3s and then the actuator (with the tethered attack target) is automatically retracted to remove the attack target. The task is self-paced, and animals are not cued or incentivized to perform the task for any other reason beyond the aggression reward itself. Poking into the “null” port is unrewarded but otherwise does not affect task timing. No food, water, or other social cues are used during training or testing. We trained 40 mice on this task for 30-45 minutes/day and 37 performed this task to criteria (defined as twice the number of pokes to the social port as null port, and a minimum of 1 poke/5 min) within 12 days (Supplementary Fig. 1b-c). We used both manual annotation and pose tracking to quantify the locations and postures of animals during both free aggression and the SOAR task^20,21^.

**Figure 1.**
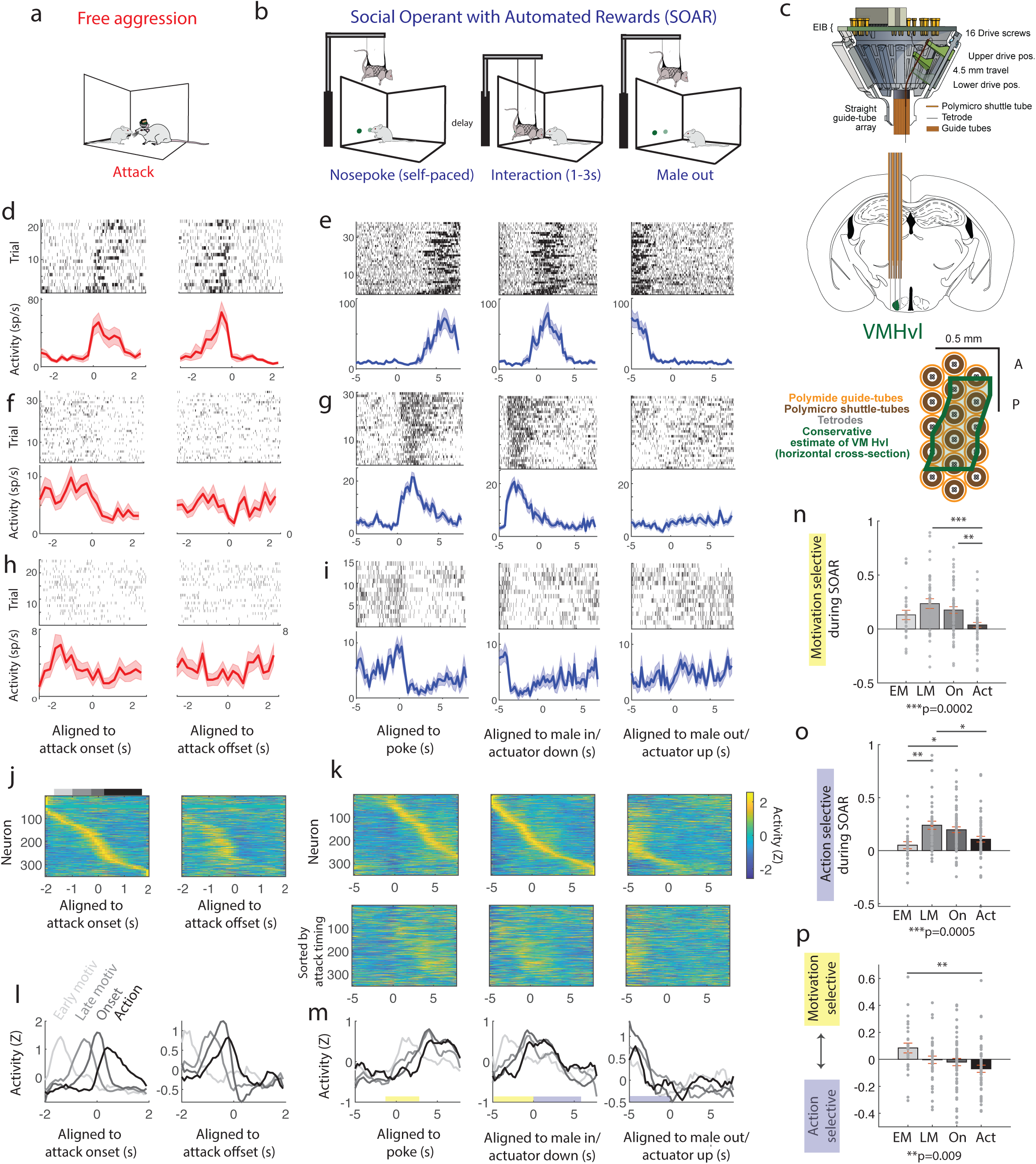
VMHvl neurons represent the sequence of motivation to action similarly across different tests of aggression. **a-b**, We recorded neurons during two types of aggression, “free” aggression in the homecage (**a**) and aggression-seeking behavior with the SOAR task which requires more stereotyped behavior during the motivational phases(**b**). **c**, We used custom shuttle drives (top) to target the full-extent of the VMHvl and its edges (middle). Shown is a reconstruction in the axial plane of the targets of the tetrodes with the outline of the VMHvl shown in green (bottom). **d-i**, Three example neurons shown in both aggression paradigms aligned to key timepoints (attack onset and offset during free aggression, red; poke onset, male in onset, and male out onset in the SOAR task, blue). Example single neuron shown in **d-e** is selective for action, while example neurons shown in **f-g** and **h-i** show examples of neurons with selectivity in the motivational phase. **j,** Population of neurons recorded in both aggression paradigms (n=353 neurons) sorted by peak time relative to attack onset. Grayscale bars (top, right) represent bins used to categorize neurons by peak response time (**l-p**). **k,** Population of neurons recorded in both aggression paradigms (n=353 neurons) sorted by peak time relative to male in onset (top) and to attack onset (bottom). Yellow and blue bars represent time bins used to compute motivation and action selectivity respectively. **l-m**, Neurons sorted into groups by their peak activation time relative to attack onset in both aggression paradigms (see gray bars, **j**, Early motivation early motivation, n=28 neurons; late motivation n=39 neurons; onset, n=59 neurons, and action, n=59 neurons). **n,** Motivation selectivity (compared to baseline) in SOAR task for classes of neurons defined by timing in free aggression (one-way ANOVA with Tukey-Kramer post hoc test for multi comparison; F_3,179_=6.778,p=0.0002 for full, late motivation vs action p=0.0001, onset vs action p=0.006). **o,** Action selectivity (compared to baseline) in SOAR task for classes of neurons defined by timing in free aggression (one-way ANOVA with Tukey-Kramer post hoc test for multi comparison; F_3,179_=6.152, p=0.0005 for full, early motivation vs late motivation p=0.0017, early motivation vs onset p=0.0138, early motivation vs action p=0.0126). **p,** Selectivity for motivation to action in SOAR task for classes of neurons defined by timing in free aggression (one-way ANOVA with Tukey-Kramer post hoc test for multi comparison; F_3,179_=4.002,p=0.009, early motivation vs action p=0.0034). For all panels *p<0.05, **p<0.01, ***p<0.001. Population responses are mean ±SEM.

### Neurons in the VMHvl region represent an aggressive motivation-to-action sequence in a behaviorally invariant manner

Though the VMHvl is spatially small, previous recordings suggest that response properties are heterogeneous^10,22^. However, it is currently unclear whether VMHvl neurons encode a motivational signal during free aggression, since it has been difficult to assay motivation in free behavior contexts. Here, to comprehensively assay the response properties of neurons in the VMHvl region, we recorded populations of neurons using custom-designed shuttle drives with spatially extended tetrode arrays (Fig.1c, n= 393 neurons across 4 animals, Supplementary Fig.1d-g) ^23^, and reconstructed the locations of recorded neurons using light-sheet microscopy (Supplementary Fig. 1f). To track neurons during aggression contexts with different patterns of motivated behaviors, we recorded the same neurons both during free aggression and the SOAR task in the same recording session.

We plotted the activity of each neuron recorded in both aggression tasks aligned to specific behavioral timepoints: the onset of attack during free aggression, the time of the nosepoke, the male in time (actuator down), and the male out time (actuator up) during the SOAR task (Fig. 1d-k). By tracking individual neurons across both aggressive behaviors, we find that individual neurons in the VMHvl region exhibit time-locked preferences for similar temporal phases of aggression during both free aggression and the SOAR task. For example, individual neurons may exhibit a peak of activity aligned to the onset of attack in free aggression and will have a peak of activity during the interaction phase of the SOAR task (Fig. 1d-e), or may exhibit peaks of activity during the phase immediately prior to attack/interaction (Fig. 1f-g), or may increase in activity during the motivational phase even earlier relative to the attack (Fig. 1h-i). Across the population, when neurons are sorted by their peak activation relative to attack during free aggression (Fig 1j) or by their peak relative to the “male-down” time (Fig 1k, top), we observe that in both cases, activity forms a temporal sequence from the motivational phase to the action phase. In addition, motivation-to-action temporal coding is preserved in the SOAR task when sorted by timing in free aggression (Fig. 1k, bottom). We classified neurons into temporal categories based on the timing of their peak activity aligned to attack onset during free aggression (Fig. 1l, early motivation, n=28 neurons; late motivation n=39 neurons; onset, n=59 neurons, and action, n=59 neurons). Using this free aggression-defined categorization, we observe that the onsets of these neurons are staggered in the same temporal order when aligned to specific timepoints during the SOAR task (Fig. 1m). In addition, we find highly significant differences in how selective these neurons are during the motivation phase of the SOAR task (Fig. 1n, relative to baseline, one-way ANOVA with Tukey-Kramer post hoc test for multi comparison; F_3,179_=6.778, p=0.0002 for full, late motivation vs action p=0.0001, onset vs action p=0.006), the action phase of the SOAR task (Fig. 1o, relative to baseline, one-way ANOVA with Tukey-Kramer post hoc test for multi comparison; F_3,179_=6.152, p=0.0005 for full, early motivation vs late motivation p=0.0017, early motivation vs onset p=0.0138, early motivation vs action p=0.0126), and how selective they are for either the motivation or action phases of the SOAR task (Fig. 1p, one-way ANOVA with Tukey-Kramer post hoc test for multi comparison; F_3,179_=4.002,p=0.009, early motivation vs action p=0.0034). Overall, we find a shift from motivation to action selectivity, with neurons with the earliest peaks during free aggression exhibiting the highest level of motivation selectivity and the neurons with the latest peaks of activity with the most action selectivity, and that this timing is conserved in both free and behaviorally-stereotyped aggression.

### Local and long-range inhibitory inputs target heterogeneous populations of VMHvl neurons

Since we observed that neurons in the VMHvl region have preferential peaks of activity during aggressive motivation or action that are preserved across aggression contexts, we sought to test whether these processes could be independently regulated by inhibitory inputs. First, we tested the hypothesis that gating would be performed by “labeled lines” that make monosynaptic contacts with subsets of neurons in the VMHvl. Here, we focused on comparing two major sources of inhibition to the VMHvl, local inhibition from the VMHvl shell, and long-range inhibition from the MPO, the VMHvl’s largest input^15^. We performed *ex vivo* whole-cell patch clamp recordings and optogenetic-assisted circuit mapping experiments (Fig. 2a). To test the connectivity of the VMHvl shell, we injected the VMHvl shell of vgat-ires-cre male mice with AAV1-Ef1-DIO-hCHR2(H134R)-mCherry-WPRE-HGHpA. In a separate cohort, to test whether VMHvl core neurons receive inhibitory input from the MPO, we injected vgat-ires-cre male mice with AAV1-Ef1-DIO-hCHR2(H134R)-eYFP-WPRE-HGHpA to the MPO and AAV2-Ef1-DIO-mCherry to the VMHvl.

**Figure 2.**
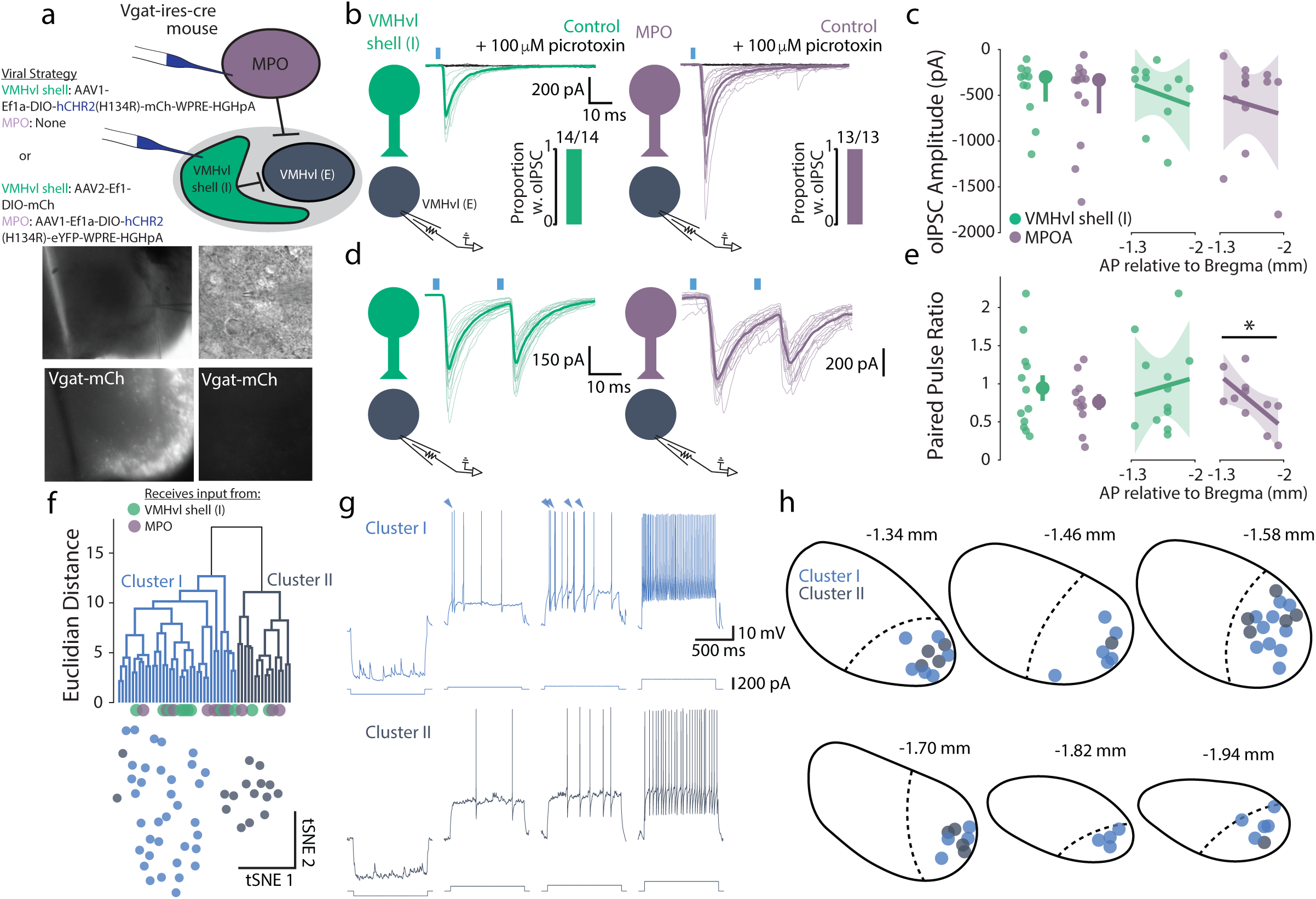
Local and long-range inhibitory inputs monosynaptically target heterogeneous VMHvl neurons. **a**, Top: Viral strategy for *ex vivo* whole cell patch clamp experiments to drive ChR2 expression in local (VMHvl shell, green, Vgat^+^) and long-range (MPO, purple, Vgat^+^) inhibitory inputs. Bottom: Example DIC and fluorescent images of a recording of a VMHvl core neuron at low and high magnification. **b**, oIPSCs recorded from representative VMHvl core neurons in response to stimulation of local (left, green) or long-range (right, purple) inhibitory inputs in control conditions (10 μM NBQX, 50 μM D-APV, in color) and following the addition of 100 μM picrotoxin (in black). Thin traces show individual trials, and thick traces show cellular mean. Insets show proportion of VMHvl core neurons receiving synaptic input from long-range or local inhibitory inputs. **c**, Left: No difference in oIPSC amplitude in VMHvl core neurons in response to stimulation of long-range or local inhibitory inputs (*p* = 0.69, Mann-Whitney U test). Right: No effect of anatomical position on VMHvl core oIPSC amplitude (local: *r*^2^ = 0.043, *F* = 0.40, *p* = 0.54, linear regression model; long-range: *r*^2^ = 0.013, *F* = 0.15, *p* = 0.71, linear regression model). **d**, Paired oIPSCs recorded from representative VMHvl core neurons in response to stimulation of local (left, green) or long-range (right, purple) inhibitory inputs. Thin traces show individual trials, and thick traces show cellular mean. **e**, Left: No difference in paired pulse ratio in VMHvl core neurons in response to stimulation of long-range or local inhibitory inputs (*p* = 0.37, unpaired t-test). Right: Anatomical position predicts paired pulse ratio in VMHvl core neuron responses to long-range inhibitory input (local: *r*^2^ = 0.012, *F* = 0.12, *p* = 0.74, linear regression model; long-range: *r*^2^ = 0.41, *F* = 6.15, *p* = 0.035, linear regression model). **f**, Hierarchical (top) and tSNE (bottom) clustering of VMHvl core neurons based on their physiological properties and anatomical position along the anterior-posterior axis of the VMHvl. Class I neurons are shown in blue and class II neurons are shown in gray. Green and purple dots below the dendrogram show the identity of neurons included in optogenetic circuit mapping experiments. **g**, Representative traces from class I (top) and class II (bottom) VMHvl core neurons. Traces show voltage responses to -100 pA, rheobase, rheobase + 20 pA, +200 pA current injections (left to right). Triangles point to action potential bursts. **h**, Anatomical position of classified VMHvl core neurons.

We do not find evidence that local and long-range inhibitory neurons target specific subsets of neurons and instead find evidence of broad connectivity. Confirming that VMHvl core neurons receive functional local and long-range inhibitory synaptic input, we find that 100% of recorded core neurons show optical inhibitory postsynaptic current (oIPSC) responses to brief blue light illumination (Fig. 2b; local: n = 14/14 neurons; long-range: n = 13/13 neurons). Confirming the GABAergic nature of the oIPSCs, application of the GABA_A_ receptor antagonist picrotoxin abolished the response (Fig. 2b). In addition, we tested whether the response of VMHvl core neurons to local or long-range inhibitory input differed and found no differences in oIPSC amplitude or paired-pulse ratio (Fig 2b-e; c, left: *p* = 0.69, Mann-Whitney U test, n_local_ = 11 neurons; long-range: 13 neurons; e, left: *p* = 0.37, unpaired t-test, n_local_ = 12 neurons, n_long-range_ = 11 neurons). As MPO-projecting VMHvl neurons are preferentially located in posterior VMHvl, we tested if VMHvl responses to inhibitory input were related to the anatomical location of the recorded neuron. Indeed, we found a significant relationship between paired-pulse ratio of long-range inputs and the position of the recorded neuron along the anterior-posterior axis of VMHvl such that the pair-pulse ratio decreased with increasing posterior position (Fig 2c,e; c, right, local: *p* = 0.54, *r*^2^ = 0.043, *F* = 0.40, linear regression model; c, right, long-range: *p* = 0.71, *r*^2^ = 0.013, *F* = 0.15, linear regression model; single stimulation n_local_ = 11 neurons; n_long-range_: 13 neurons; e, right, local: *p* = 0.74, *r*^2^ = 0.012, *F* = 0.12, linear regression model; e, right, long-range: *p* = 0.035, *r*^2^ = 0.41, *F* = 6.15, linear regression model; paired pulse stimulation n_local_ = 12 neurons, n_long-range_ = 11 neurons). These data suggest that while both local and long-range inhibitory inputs to VMHvl broadly target core neurons, long-range inhibitory MPO inputs show greater presynaptic strength at posterior VMHvl sites.

We also tested whether inhibitory inputs had preferential contact with subclasses of neurons defined by their functional properties. Prior work has suggested that VMHvl core neurons are a heterogeneous population of primarily glutamatergic neurons with diverse gene expression, projection patterns, firing properties, and behavioral function^10,15,19,24,25^. Indeed, we find that VMHvl core neurons can be divided into two populations on the basis of their anterior-posterior position and physiological properties (Fig 2f-h; see Methods for details on clustering procedures). We find that approximately 70% of the neurons can be sorted into cluster I whereas the remainder fall into cluster II, and that both populations are found across the anterior-posterior extent of the VMHvl (Fig. 2h; *p* = 0.66, Mann-Whitney U test, n_cluster I_ = 31 neurons, n_cluster II_ = 16 neurons). Cluster I and II neurons show a variety of differences in their physiological properties (Fig. 2g; Table 1; n_cluster I_ = 31 neurons, n_cluster II_ = 16 neurons). Most notably, compared to cluster II neurons, cluster I neurons display a greater maximum firing rate (*p* = 3.36 × 10^-4^, unpaired t-test), a bias towards firing action potentials earlier in response to a sustained current injection (firing bias: *p* = 0.037, Mann-Whitney U test; first action potential latency: *p* = 0.0049, Mann-Whitney U test), a propensity to fire bursts of action potentials at the start of a train of action potentials (first ISI: 2.12 × 10^-4^, Mann-Whitney U test; second ISI: 1.11 × 10^-5^, Mann-Whitney U test), and greater amplitude accommodation (*p* = 0.0028, Mann-Whitney U test) as well as action potential halfwidth broadening (*p* = 2.86 × 10^-4^, Mann-Whitney U test) during sustained firing. Cluster II neurons, in comparison to cluster I neurons, are characterized by their fast, narrow action potentials exemplified by their action potential slope (Rising slope: *p* = 0.022, unpaired t-test; Downslope: *p* = 4.26 × 10^-4^, unpaired t-test) action potential halfwidth (*p* = 0.014, Mann-Whitney U test), and after-hyperpolarization potential (Latency: *p* = 8.39 × 10^-4^, Mann-Whitney U test; Rise: *p* = 3.57 × 10^-4^, unpaired t-test).Overall, we find that neurons from both clusters are targeted by local and long-range inhibitory input, indicating that these inputs can broadly regulate the activity of the heterogeneous VMHvl core, though the weight of the input from the MPO may be strongest in the posterior VMHvl.

**Table 1.**
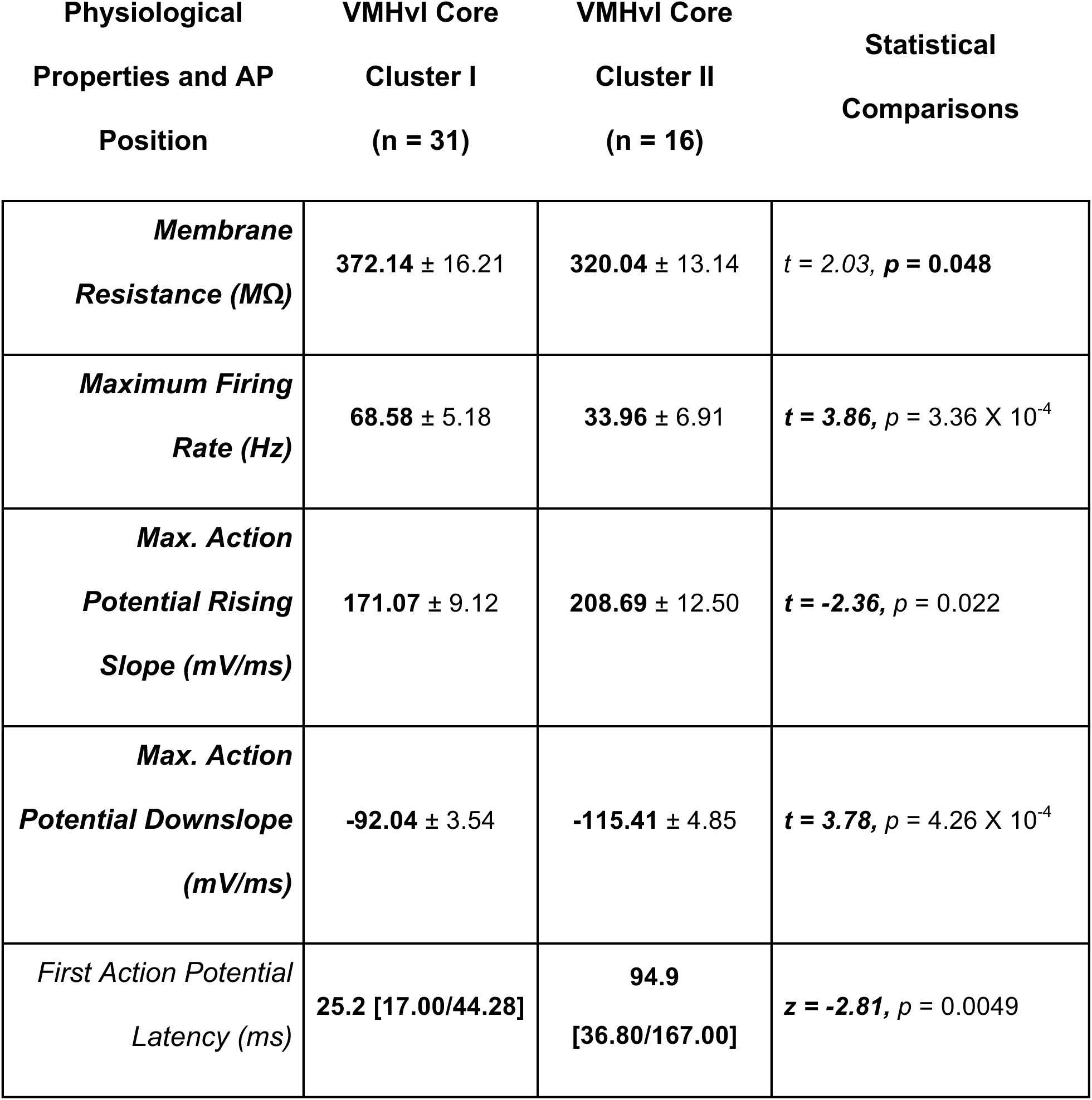
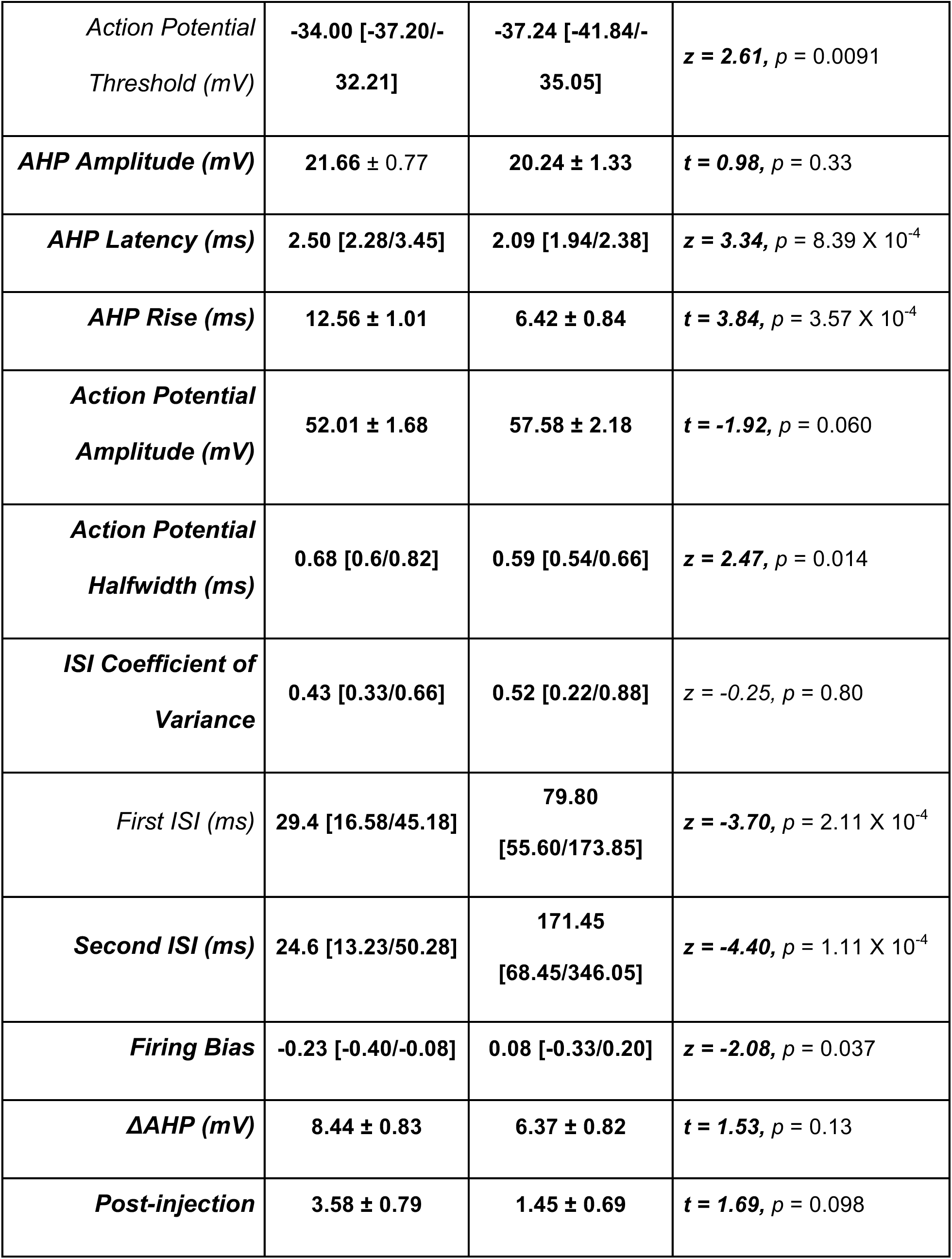
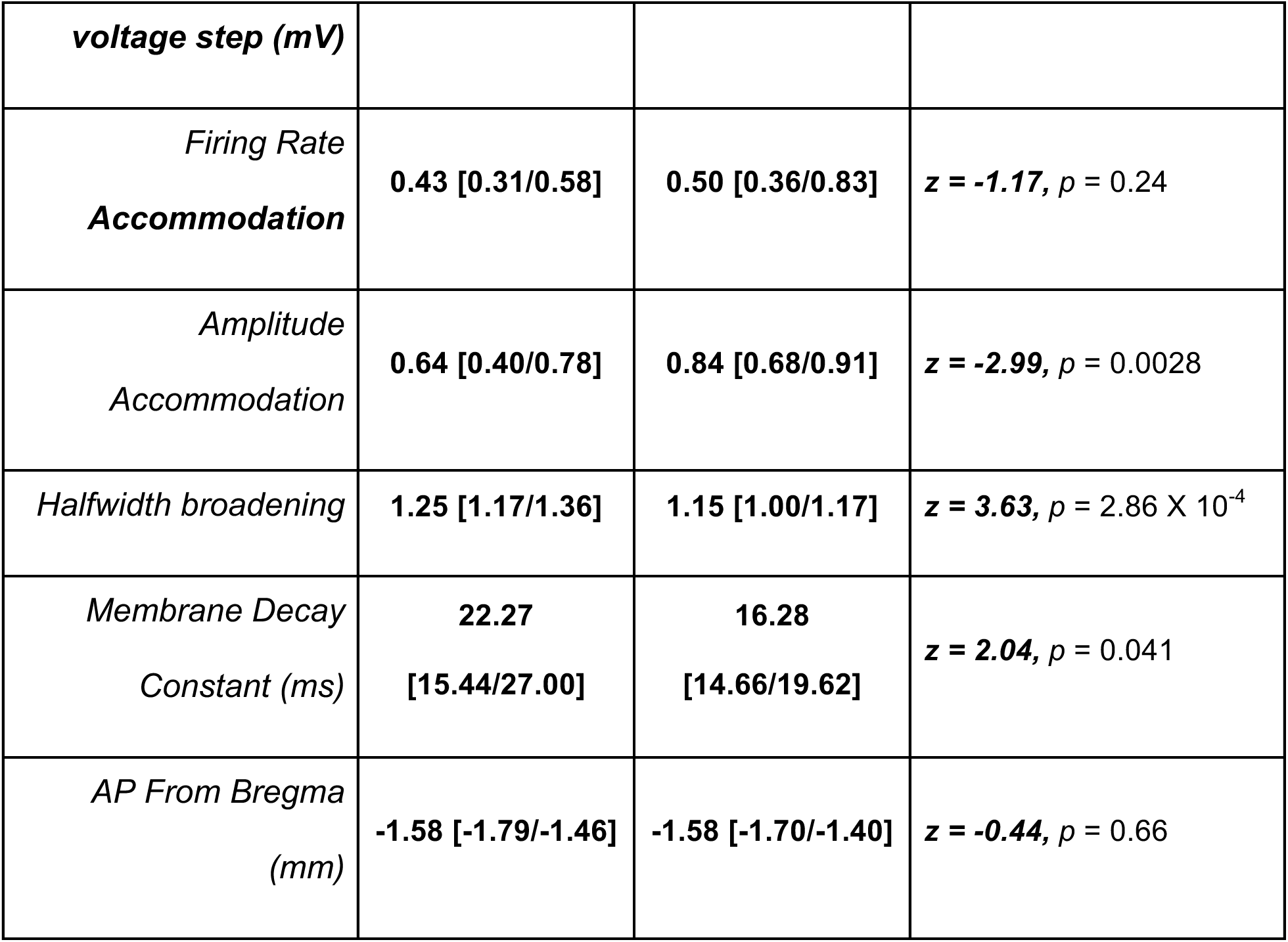
Physiological properties and anterior posterior position of VMHvl core neuron clusters I and II. Normal data presented as mean ± s.e.m. with differences tested using an unpaired t-test. Non-normal data are presented as median and interquartile range [1st quartile/3rd quartile] with differences tested using a Mann-Whitney U test. Significant comparisons bolded.

**Table 2.**
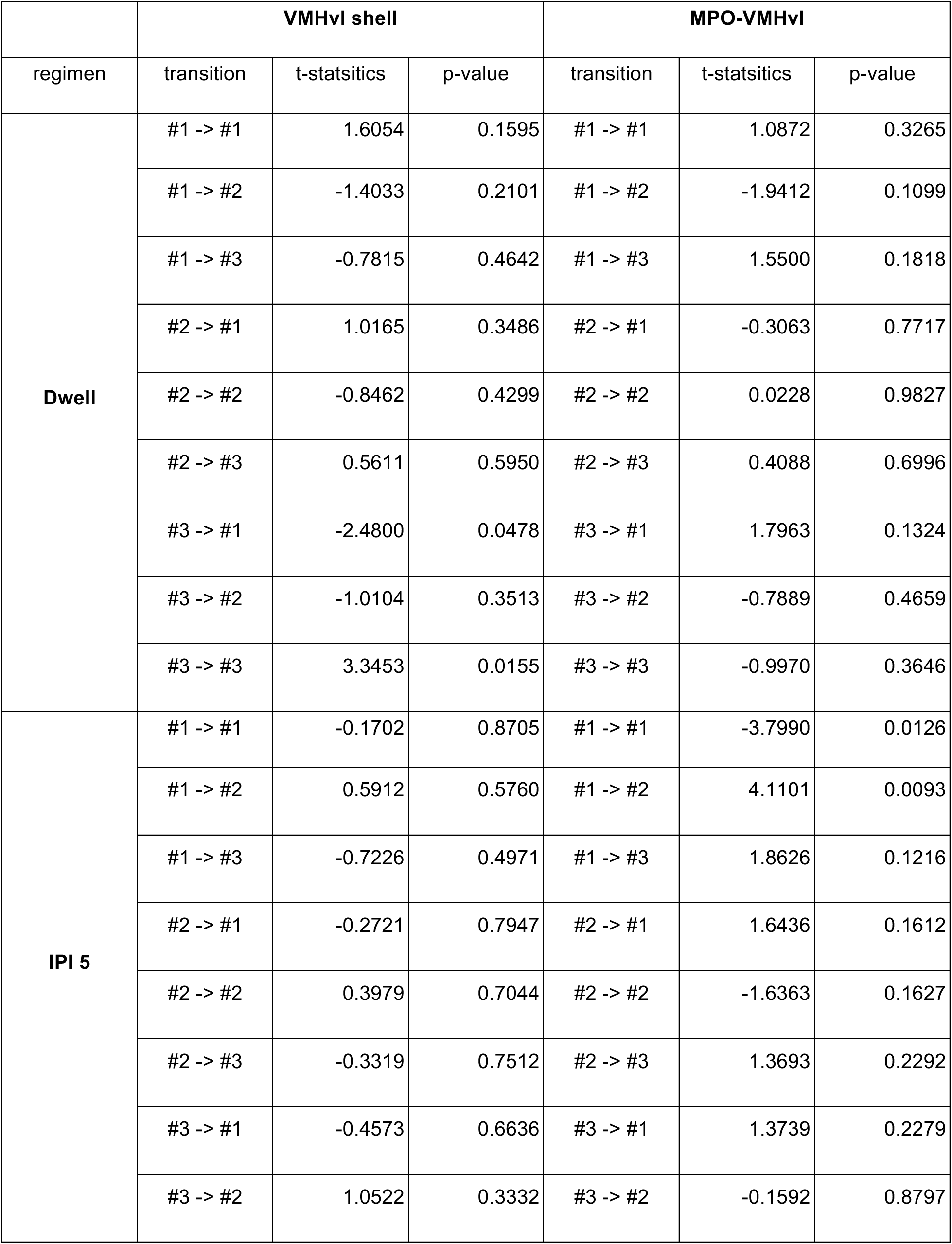
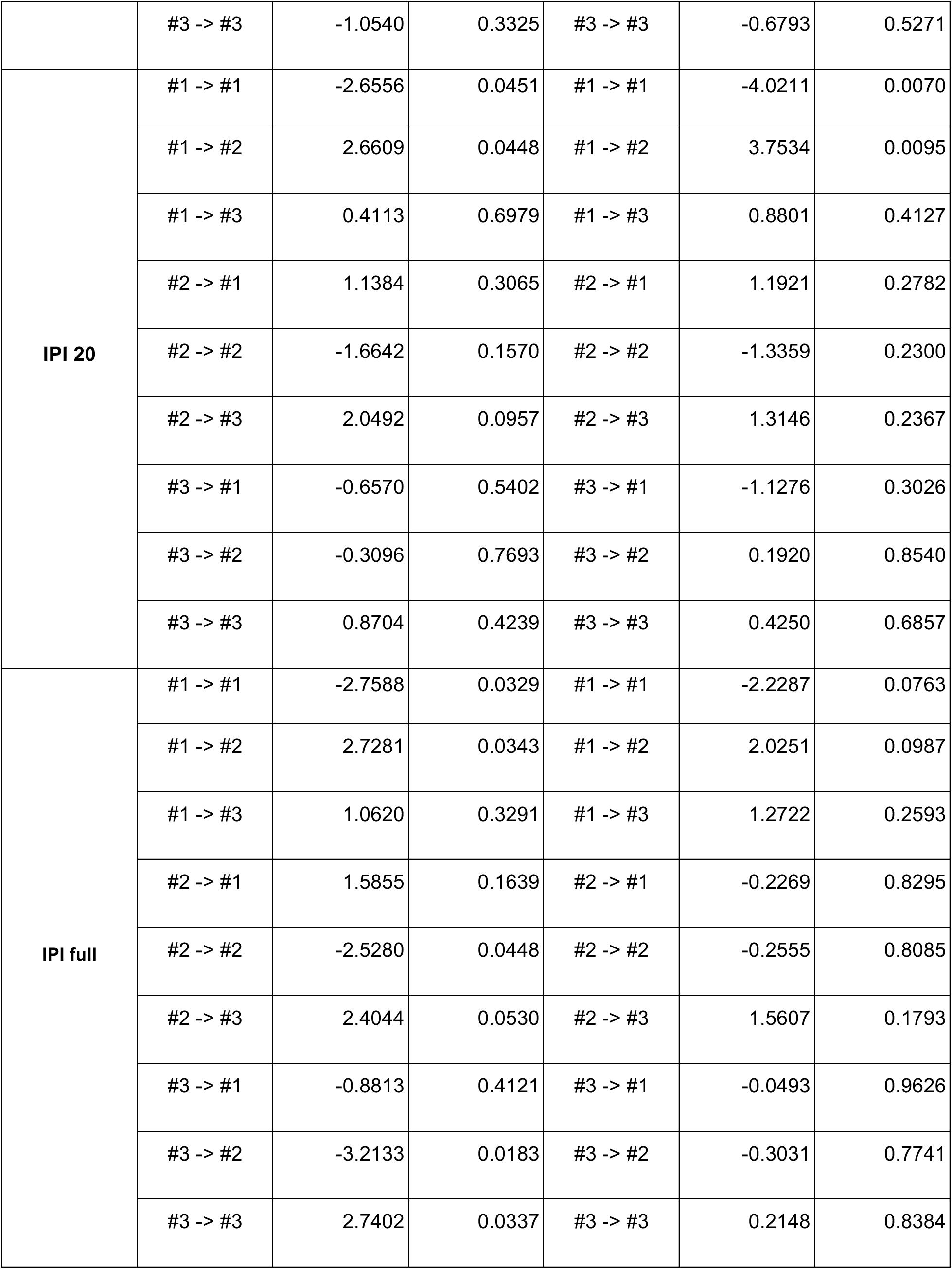
Comparisons of the state-transition probabilities between sham and stim trials. Paired t-test was performed for each comparison.

|  | VMHvl shell |  |  | MPO-VMHvl |  |  |
| --- | --- | --- | --- | --- | --- | --- |
| regimen | transition | t-statsitics | p-value | transition | t-statsitics | p-value |
| **Dwell** | #1 -> #1 | 1.6054 | 0.1595 | #1 -> #1 | 1.0872 | 0.3265 |
|  | #1 -> #2 | -1.4033 | 0.2101 | #1 -> #2 | -1.9412 | 0.1099 |
|  | #1 -> #3 | -0.7815 | 0.4642 | #1 -> #3 | 1.5500 | 0.1818 |
|  | #2 -> #1 | 1.0165 | 0.3486 | #2 -> #1 | -0.3063 | 0.7717 |
|  | #2 -> #2 | -0.8462 | 0.4299 | #2 -> #2 | 0.0228 | 0.9827 |
|  | #2 -> #3 | 0.5611 | 0.5950 | #2 -> #3 | 0.4088 | 0.6996 |
|  | #3 -> #1 | -2.4800 | 0.0478 | #3 -> #1 | 1.7963 | 0.1324 |
|  | #3 -> #2 | -1.0104 | 0.3513 | #3 -> #2 | -0.7889 | 0.4659 |
|  | #3 -> #3 | 3.3453 | 0.0155 | #3 -> #3 | -0.9970 | 0.3646 |
| **IPI 5** | #1 -> #1 | -0.1702 | 0.8705 | #1 -> #1 | -3.7990 | 0.0126 |
|  | #1 -> #2 | 0.5912 | 0.5760 | #1 -> #2 | 4.1101 | 0.0093 |
|  | #1 -> #3 | -0.7226 | 0.4971 | #1 -> #3 | 1.8626 | 0.1216 |
|  | #2 -> #1 | -0.2721 | 0.7947 | #2 -> #1 | 1.6436 | 0.1612 |
|  | #2 -> #2 | 0.3979 | 0.7044 | #2 -> #2 | -1.6363 | 0.1627 |
|  | #2 -> #3 | -0.3319 | 0.7512 | #2 -> #3 | 1.3693 | 0.2292 |
|  | #3 -> #1 | -0.4573 | 0.6636 | #3 -> #1 | 1.3739 | 0.2279 |
|  | #3 -> #2 | 1.0522 | 0.3332 | #3 -> #2 | -0.1592 | 0.8797 |
|  | #3 -> #3 | -1.0540 | 0.3325 | #3 -> #3 | -0.6793 | 0.5271 |
| IPI 20 | #1 -> #1 | -2.6556 | 0.0451 | #1 -> #1 | -4.0211 | 0.0070 |
|  | #1 -> #2 | 2.6609 | 0.0448 | #1 -> #2 | 3.7534 | 0.0095 |
|  | #1 -> #3 | 0.4113 | 0.6979 | #1 -> #3 | 0.8801 | 0.4127 |
|  | #2 -> #1 | 1.1384 | 0.3065 | #2 -> #1 | 1.1921 | 0.2782 |
|  | #2 -> #2 | -1.6642 | 0.1570 | #2 -> #2 | -1.3359 | 0.2300 |
|  | #2 -> #3 | 2.0492 | 0.0957 | #2 -> #3 | 1.3146 | 0.2367 |
|  | #3 -> #1 | -0.6570 | 0.5402 | #3 -> #1 | -1.1276 | 0.3026 |
|  | #3 -> #2 | -0.3096 | 0.7693 | #3 -> #2 | 0.1920 | 0.8540 |
|  | #3 -> #3 | 0.8704 | 0.4239 | #3 -> #3 | 0.4250 | 0.6857 |
| IPI full | #1 -> #1 | -2.7588 | 0.0329 | #1 -> #1 | -2.2287 | 0.0763 |
|  | #1 -> #2 | 2.7281 | 0.0343 | #1 -> #2 | 2.0251 | 0.0987 |
|  | #1 -> #3 | 1.0620 | 0.3291 | #1 -> #3 | 1.2722 | 0.2593 |
|  | #2 -> #1 | 1.5855 | 0.1639 | #2 -> #1 | -0.2269 | 0.8295 |
|  | #2 -> #2 | -2.5280 | 0.0448 | #2 -> #2 | -0.2555 | 0.8085 |
|  | #2 -> #3 | 2.4044 | 0.0530 | #2 -> #3 | 1.5607 | 0.1793 |
|  | #3 -> #1 | -0.8813 | 0.4121 | #3 -> #1 | -0.0493 | 0.9626 |
|  | #3 -> #2 | -3.2133 | 0.0183 | #3 -> #2 | -0.3031 | 0.7741 |
|  | #3 -> #3 | 2.7402 | 0.0337 | #3 -> #3 | 0.2148 | 0.8384 |

### Local inhibition and VMHvl neurons modulated by long-range inhibitory inputs show distinct motivation and action signatures

While we observed in *ex vivo* recordings that both local and long-range inhibitory inputs target VMHvl core neurons broadly, (i.e. we observed that 100% of neurons we recorded from in each case exhibited light-evoked IPSCS, Fig. 2b), we found that the strength of the synaptic input varies across the anterior-posterior axis of the VMHvl for the MPO but not the VMHvl shell. This result suggests that specificity of function is not performed by targeting separable subpopulations, and instead, differences in control may be achieved through preferred functional connectivity to post-synaptic partners or through gating at specific timepoints during the motivation-action sequence.

To address these possibilities, we used optogenetic perturbation to perform “identity tagging” of neurons, allowing us to identify neurons with cell-type specificity or neurons that are modulated by optogenetic perturbation of these populations. At the time of electrode implantation, vgat-ires-cre mice were injected with AAV2-Ef1-DIO-hCHR2(H134R) in the VMHvl shell and AAV2-Ef1a-DIO-eNpHR3.0 in the MPO and stimulating fibers (200 μm) were placed over the VMHvl shell and MPO (Supplementary Fig 1g). Optogenetic perturbation was always performed at the end of each behavioral session so it would not influence recorded behaviors. After recording behavior for each session including both the SOAR task and free aggression, we looked for specific patterns of light-evoked modulation, allowing us to identify units as GABAergic VMHvl shell neurons, neurons suppressed by VMHvl shell, and neurons disinhibited by MPO (Fig 3a-g). Specifically, we delivered a series of brief pulses of blue light (460nm, 5ms, 1-4Hz) in order to optically identify VMHvl shell^vgat+^ neurons and to detect VMHvl core neurons that received short latency inhibition from this input. To detect VMHvl neurons modulated by MPO (looking specifically for neurons that are disinhibited by MPO^vgat+^ neurons), we delivered a series of pulses of yellow light (556nm, 500ms, 0.1-0.2Hz). We identified neurons of each class (VMHvl shell optotagged, VMHvl-shell suppressed, and MPO modulated) by comparing the activity during the laser to a baseline no laser period for each neuron (Fig. 3e-g, significance corrected for multiple comparisons using Benjamini-Hochberg correction). We also detected a small number of VMHvl neurons suppressed by MPO inhibition (Supplementary Fig 3a-c). We then compared these identity tags to the response profiles of the neurons during motivation and action behaviors recorded during free aggression and during the SOAR task.

**Figure 3.**
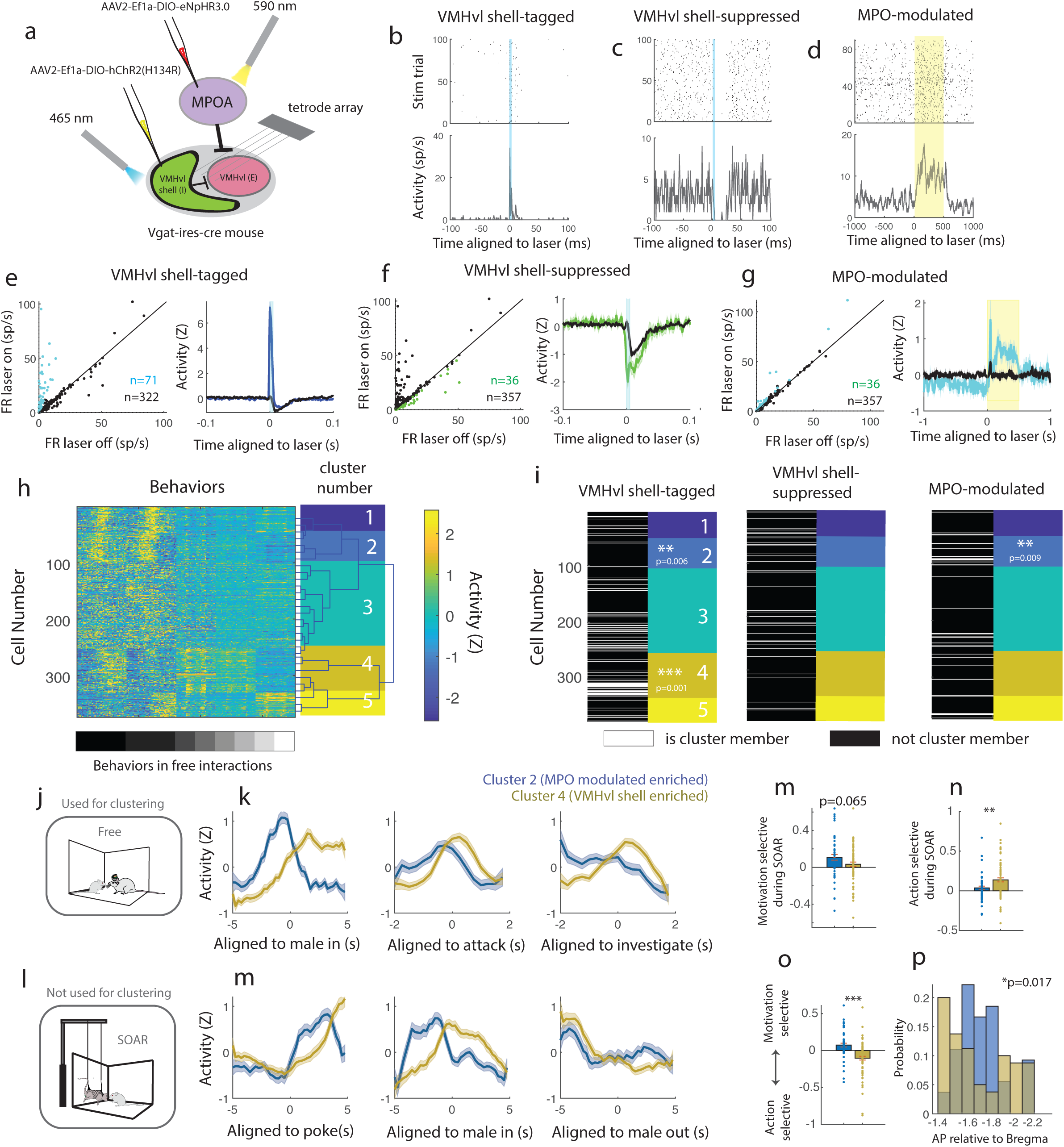
Optogenetic identity-tagged neurons show distinct motivation and action selectivity. **a,** Viral and recording strategy for optogenetic identity-tagging experiments. **b-d,** Rasters and PSTHs for example identity-tagged neurons for VMHvl shell optotagged (**b,** vgat+ neurons in the VMHvl shell), VMHvl shell suppressed (**c,** VMHvl neurons suppressed by activation of the VMHvl shell^vgat+^), and MPO-modulated through disinhibition (**d,** VMHvl neurons disinhibited by inactivation of MPO^vgat+^). Color bar indicates duration of light pulse for identity-tagging. **e-g,** Comparison of laser-evoked and baseline responses (left) and population laser aligned PSTH (right) for each class of identity-tagged neurons, VMHvl shell optotagged (**e**), VMHvl shell-suppressed (**f**) and MPO modulated (**g**). n=393 neurons for experiments. See methods for identity tagging criteria, all comparisons corrected with false discovery rate correction. **h,** Activity matrix and unsupervised clustering of neural dynamics aligned to behaviors during free interaction. Behaviors on the x-axis from left to right marked by grayscale bar are introduction of the male, introduction of the female, attack onset, bite, punch, tail rattle, investigation of male, investigation of female. **i,** Distribution of cluster for each class of identity-tagged neurons (cluster sorting and color coding is the same as in **h**). White bars indicate that the neurons located in that row is an identity-tagged neuron for VMHvl shell optotagged (left), VMHvl shell-suppressed (center), and MPO modulated (right). Cluster identity is shown to the right of each distribution of identity-tagged neurons using the colors from **h**. VMHvl shell-tagged neurons are enriched in cluster 4 (p=0.001, Fisher’s exact test, Bonferroni correction), and are excluded from cluster 2 (p=0.006), while MPO-modulated identity-tagged neurons are enriched in cluster 2 (p=0.009). **j-k,** Comparison of the response properties of cluster 2 (n=54 neurons, blue) and cluster 4 (n=80 neurons, gold) of activity used for clustering, aligned to male in during free aggression (**k,** left), attack onset during free aggression (**k**, middle), and male investigate during free aggression (**k**, right). **l-m,** Comparison of the response properties on held-out data not used for clustering, SOAR task aligned to specific behavior timepoints. In both cases, activity in cluster 2 (MPO-modulated enriched) is activity earlier than cluster 4. **n-q,** Statistical comparison of the response properties and anatomical position from the held-out data from the SOAR task. Motivational selectivity (**n**, t(122)=-1.858, p= 0.170, unpaired t-test), Action selectivity (**o**, t(122)=2.696, p=0.009, unpaired t-test), Motivation vs Action selectivity (**p,** unpaired t-test, t(122)=-4.493, p=0.00002), anatomical position of neuron on the A-P axis (**q,** p=0.017, Kolmogorov-Smirnov test). For all panels *p<0.05, **p<0.01, ***p<0.001. Plots show mean±SEM.

We used unsupervised hierarchical clustering to test whether identity-tagged neurons had similar response properties and would be “enriched” in specific clusters. Since we reasoned that the timing of the response properties relative to key behaviors is the key property to delineate motivation and action related neurons, we clustered a matrix of the concatenated dynamics aligned at specific behavioral events during free interactions with males and females (introduction of the male, introduction of the female, attack onset, bite, punch, tail rattle, investigation of male, investigation of female) and found that these neural responses grouped into 5 main clusters (Fig. 3h). We did not include responses from the SOAR task in the clustering and instead used these neural responses as “held-out” data. We found that identified VMHvl shell^vgat+^ optotagged neurons (n=71 neurons) were significantly more likely to belong to cluster 4 than predicted by chance (Fig. 3i, p=0.0011, Fisher’s exact test, with Bonferroni correction for tests on multiple clusters). In addition, VMHvl shell optotagged neurons were significantly excluded from cluster 2 (p=0.006, Fisher’s exact test, with Bonferroni correction for multiple clusters). In contrast, MPO modulated neurons, though rarer (n=36 neurons) were more likely than chance to belong to cluster 2 (p=0.0088, Fisher’s exact test, with Bonferroni correction for multiple clusters). VMHvl shell-suppressed neurons were not more likely to belong to any particular cluster (p>0.05 for all clusters, Fisher’s exact test), though loosening our stringent inclusion criteria revealed a preference for cluster 1 (Supplementary Fig. 2d-f).

Next, we examined the response properties of clusters identified as being enriched in our identity-tagged units. We found that the activity profiles from cluster 2 (enriched in MPO-modulated units) were leftward shifted in time relative to cluster 4 (n=71 enriched in VMHvl shell^vgat+^ optotagged neurons), suggesting that these groups of neurons were preferentially active during aggressive motivation or action respectively (Fig. 3j-k). Surprisingly, we found the same patterns of activation when we compared the response profiles of these enriched clusters aligned to behaviors in the SOAR task (held-out data). We observed that neurons in the MPO-modulated enriched cluster (n=36, cluster 2, Fig 3h), were more active when aligned to the nosepoke prior to the start of the interaction phase of the SOAR task (Fig. 3j) than the VMHvl shell enriched cluster (cluster 4). Additionally, neurons in the MPO-modulated enriched cluster were significantly more motivation selective compared to action selective (Fig. 3n-p, action selectivity p=0.0086, and p=0.00006 for motivation vs action selectivity, paired t-test) and were anatomically more biased toward the posterior part of the A-P axis (Fig. 3q, p=0.0017 Kolmogorov-Smirnov test). Together, these data extend and are consistent with the results of our circuit-mapping experiments, showing that VMHvl shell^vgat+^ neurons are action selective and that VMHvl shell-suppressed units have heterogeneous response characteristics. In contrast, neurons modulated by MPO inhibition are motivation-selective than action selective, and may have preferential coupling with neurons in the posterior VMHvl (Fig. 2e,3q).

### Local and long-range inhibitory populations have distinct activity signatures during aggressive motivational and action epochs

Inhibitory inputs to the VMHvl core may gate output during distinct phases of the motivation-action sequence. Thus, we next directly compared the differences in the response selectivity and temporal dynamics during aggression between the local and long-range inhibitory inputs during aggressive motivation and action. To do this, we recorded the population responses of local inhibitory neurons in the VMHvl shell and long-range inhibitory MPO-VMHvl neurons using fiber photometry during free social interactions and during the SOAR task. To record from local inputs, we injected vgat-ires-cre males with AAV2-Syn-DIO-GCaMP6f in the VMHvl shell and implanted a fiber over the VMHvl core (Fig. 4a, Supplementary Fig. 3a). Since the VMHvl shell and VMHvl core are anatomically very close, we are likely recording from cell soma as well as terminals, so will refer to this population as VMHvl shell^vgat+^. To record from long-range inputs, we injected a retrogradely transported calcium indicator (AAVrg-Ef1a-DIO-GCaMP6f) into the VMHvl core of vgat-ires-cre male and implanted a recording fiber over the MPO (Fig 4b, Supplementary Fig. 3b). Control animals were injected with eGFP (AAV5-EF1a-eGFP,, Supplementary Fig. 3c-f). We performed fiber photometry recordings from VMHvl shell^vgat+^ and MPO-VMHvl^vgat+^ populations not only during free aggression and the SOAR task, but also during free interactions with females and pups (Fig. 4c). All males used in this recording were sexually and parentally experienced.

**Figure 4.**
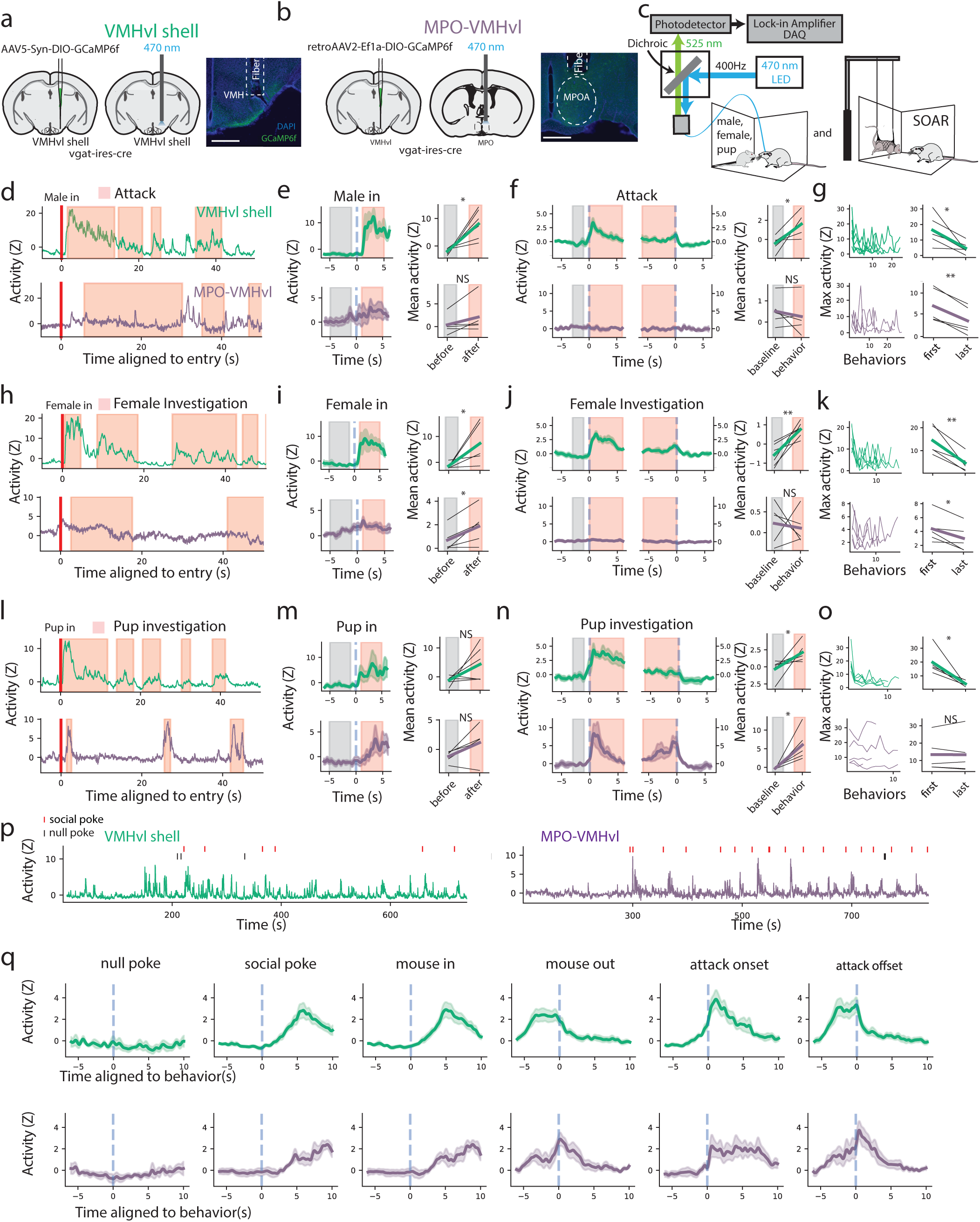
Local and long-range inhibitory inputs to the VMHvl have unique activity profiles during aggressive motivation, aggressive action, and other social behaviors. **a,** Viral strategy and a histological image showing the location of the optic fibers to record GCaMP6f activity in the VMHvl shell^vgat+^ neurons. **b**, Viral strategy and a histological image showing the location of the optic fibers to record GCaMP6f activity in the MPO-VMHvl^vgat+^ neurons. For (**a**) and (**b**), scale bars (white), 500 µm. Fiber tips, 400 µm diameter. **c,** We recorded neural activity during free aggression and interactions with females and pups (left) and during the SOAR task (right). **d,** Example traces of the z-score of GCaMP6f activities in the VMHvl shell^vgat+^ (green, top) and MPO-VMHvl^vgat+^ (purple, bottom) during free aggression. The red vertical lines show the entry of male and the red indicates attack. **e,** Left, population averaged z-scored GCaMP6f activities in the VMHvl shell^vgat+^ (green, top) and MPO-VMHvl^vgat+^ (purple, bottom) aligned to the male intruder entries. Right, comparisons of AUCs of z-scored GCaMP6f activities before (−5s to -1s, gray) versus after (1s to 5s, red) male entry. Entries were significantly higher for the VMHvl shell^vgat+^ (t(5) = −3.825, p = 0.012, n = 6, paired t-test) but not for the MPO-VMHvl^vgat+^ (t(4) =-1.719, p = 0.161, n = 5, paired t-test). **f,** Left, average z-scored GCaMP6f activities in the VMHvl shell^vgat+^ (green, top) and MPO-VMHvl^vgat+^(purple, bottom) aligned to onsets and offsets of attack. Right, comparisons of AUCs of z-scored GCaMP6f activities for the baseline (−3.5s to −0.5s) versus the behavior windows (red). Attack activity was significantly higher compared to the baseline for the VMHvl shell^vgat+^ (t(5) = −3.590, p = 0.016, n = 6, paired t-test) but not for the MPO-VMHvl^vgat+^ (t(4) = 0.950, p = 0.396, n = 5, paired t-test). **g,** Signal decreases across the session. Maximum value of the signal during each bout of attack from the first to the last bout within a free aggression session. The first attack against male was significantly higher than the one in the last attack for both the VMHvl shell^vgat+^ (t(5) = 3.163, p = 0.025, n = 6, paired t-test) and the MPO-VMHvl^vgat+^ (t(4) = 5.949, p = 0.004, n = 5, paired t-test). **h.** Example traces of the z-scores of GCaMP6f activities in the VMHvl shell^vgat+^ (green) and MPO-VMHvl^vgat+^ (purple) during interaction with a female. The red vertical lines show the entries of the female and the shadings show investigation towards the female. **i.** Same as **e**, but for female entry. Activity was increase for both the VMHvl shell^vgat+^ (t(5) = − 2.790, p = 0.038, n = 6, paired t-test) and the MPO-VMHvl^vgat+^ (t(4) = 2.989, p = 0.040, n = 5, paired t-test). **j.** Same as **f**, but for investigation of a female. Activity was significantly higher compared to the baseline for the VMHvl shell^vgat+^ (t(5) = 4.235, p = 0.008, n = 6, paired t-test) but not for the MPO-VMHvl^vgat+^ (t(4) = 0.341, p = 0.750, n = 5, paired t-test). **k.** Same as **g**, but for investigation of a female. Activity was significantly higher than the one in the last investigation for both the VMHvl shell^vgat+^ (t(5) = 5.195, p = 0.0035, n = 6, paired t-test) and the MPO-VMHvl^vgat+^ (t(4) = 2,938, p = 0.042, n = 5, paired t-test). **l.** Example traces of the z-score of GCaMP6f activity in the VMHvl shell^vgat+^ (green, top) and MPO-VMHvl^vgat+^ (purple) during interaction with a novel pup. The red vertical lines show the entries of the pup and the shadings show investigation towards the pup. **m.** Same as **e**, but for pup entry. Activity was significantly higher following pup entry for the VMHvl shell^vgat+^ (t(5) = 1.898, p = 0.116, n = 6, paired t-test) and the MPO-VMHvl^vgat+^ (t(4) = 2.310, p = 0.082, n = 5, paired t-test). **n.** Same as **f**, but for pup investigation. Activity during pup investigation was significantly higher compared to the baseline for both the VMHvl shell^vgat+^ (t(5) = −2.625, p = 0.047, n = 6, paired t-test) and for the MPO-VMHvl^vgat+^ (t(4) = 3.173, p = 0.034, n = 5, paired t-test). **o.** Same as **g**, but for pup investigation. Activity was significantly higher than the first investigation for the VMHvl shell^vgat+^ (t(5) = 4.190, p = 0.009, n = 6, paired t-test) but not for the MPO-VMHvl^vgat+^ (t(4) = 0.03835, p = 0.971, n = 5, paired t-test). **p,** Example traces of the z-scores of GCaMP6f activities in the VMHvl shell^vgat+^ (green) and MPO-VMHvl^vgat+^ (purple) during SOAR task sessions. The red and black ticks above the traces show the social and null pokes during the sessions, respectively. **q,** Averaged GCaMP6f activity in the VMHvl shell^vgat+^ (top row, green) and MPO-VMHvl^vgat+^ (bottom row, purple) aligned to the different events in the SOAR task. Solid lines show mean across mice and shaded regions show standard error. From left to right, each column corresponds to null pokes, social pokes, mouse in, mouse out, attack onset, and attack offset. For all panels *p<0.05, **p<0.01, ***p<0.001. Plots show mean±SEM.

We examined the average activity aligned to specific behavior timepoints during free interactions with social targets and compared the mean signal during the behavior to the baseline preceding each behavior. During free aggression, activity in the VMHvl shell^vgat+^ population increased significantly during the introduction of males and during attack (Fig. 4d-f, top, green; entry of males: t(5) = −3.825, p = 0.012, n = 6; attack: t(5) = −3.590, p = 0.016, n = 6; paired t-test). The MPO-VMHvl^vgat+^ population response was not significantly modulated by either the entrance of the male or attack (Figure 4d, bottom, purple; entry of males: t(4) =1.719, p = 0.161, n = 5; attack: t(4) = 0.950, p = 0.396, n = 5; paired t-test), and instead, the signal appeared to increase between attacks. Fleeing events (defined by the intruder moving quickly away from the resident) were rarer and did not occur in every session, but robustly increased the signal for MPO-VMHvl^vgat+^ population (but not the VMHvl shell^vgat+^ population) when they were coincident with the end of attack (Supplementary Fig. 3a-d).

We also compared the response during non-aggressive social behaviors. The VMHvl shell^vgat+^ population increased activity at the arrival of the female and also increased robustly during investigation of the female (Fig. 4h-j, top, green; entry of females: t(5) = −2.790, p = 0.038, n = 6; female investigation: t(5) = 4.235, p = 0.008, n = 6; paired t-test). The MPO-VMHvl^vgat+^ population response modestly increased following the entrance of the female but was not affected by the action of investigation (Fig. 4h-j, bottom, purple; entry of females: t(4) = 2.989, p = 0.040, n = 5; female investigation: t(4) = 0.341, p = 0.750, n = 5; paired t-test). Surprisingly, interactions with a pup strongly and consistently activated both VMHvl shell^vgat+^ and MPO-VMHvl^vgat+^ population responses and activity decreased acutely at the offset of interactions (Fig. 4n; VMHvl shell^vgat+^: t(5) = −2.625, p = 0.047; MPO-VMHvl^vgat+^: t(4) = 3.173, p = 0.034, n = 5; paired t-test). While other interactions that drove increases in activity had strong “first trial” effects (i.e., the activity peak was highest during the first instance of the behavior, Fig. 4g,k,o), we did not observe this effect for the MPO-VMHvl^vgat+^ population response to pups, suggesting that each instance of interaction activated the MPO-VMHvl^vgat+^ population to a similar extent (Fig. 4o, bottom) (VMHvl shell^vgat+^: t(5) = 3.163, p = 0.025 for attack, t(5) = 5.195, p = 0.0035 for female investigation, t(5) = 4.190, p = 0.009 for pup investigation, n = 6; MPO-VMHvl^vgat+^: t(4) = 5.949, p = 0.004 for attack, t(4) = 2,938, p = 0.042 for female investigation, t(4) = 0.03835, p = 0.971 for pup investigation, n = 5; paired t-test). Together these data suggest that with the notable exception of pup investigation, the VMHvl shell^vgat+^ population is strongly driven by actions, and the MPO-VMHvl^vgat+^ population is activated at timepoints where aggression would be inappropriate, suggesting a role for the MPO in preventing the initiation of aggressive motivation or exiting an aggressively motivated state.

We next examined the time-locked responses of both the VMHvl shell^vgat+^ and MPO-VMHvl^vgat+^ populations during the SOAR task, where motivation and action can be more clearly assessed. We observed that trials where the resident poked the social port were associated with reliably increased activity timed to each trial in both populations (Fig. 4p). Activity in the VMHvl shell^vgat+^ population was timed to the onset of attack (Fig 4q, top), while peaks in the MPO-VMHvl^vgat+^ population were timed to the offset of attack (Fig 4q, bottom). We did not observe a significant time-locked difference in activity in GFP controls in either the free interactions or the SOAR task (Supplementary Fig.3e-h; t(2) = −0.220, p = 0.846 for male entries; t(2) = 0.336, p = 0.769 for female entries; t(2) = −1.095, p = 0.387 for attack; t(2) = −0.610, p = 0.604 for female investigation; n = 3, paired t-test).

Since behaviors in both the free aggression and SOAR task can be fast and overlapping, we used an encoding model to segregate the contributions of different temporally adjacent behavior events to the signal^26^. We modeled the signal from each mouse for both the free aggression and the SOAR task (Fig. 5a). Each model uses as inputs a series of discrete events convolved with a spine basis and also included a set of continuous predictors that we hypothesized might have additional contributions to the signal (Fig. 5b). We determined the best fit model for each mouse and examined the time-varying kernels aligned to discrete timepoints for the VMHvl shell^vgat+^ population and the MPO-VMHvl^vgat+^ population (Fig. 5c-f). Using this more principled approach to define the relationship between ongoing behavior and neural signal, we found that during free behavior, the VMHvl shell^vgat+^ population (but not the MPO-VMHvl^vgat+^ population) is strongly modulated at the onset of attack. In contrast, the MPO-VMHvl^vgat+^ population (but not the VMHvl shell^vgat+^ population) is acutely activated when the intruder flees (Fig. 5e, t(5) = − 3.83, p = 0.012, n = 6 for the kernel for attack onsets for VMHvlshell^vgat+^; t(4) = −3.395, p = 0.0426, n = 5 for the kernel for intruder fleeing for MPO-VMHvl^vgat+^). In the SOAR task, we find that the action of the nosepoke itself did not robustly modulate either signal (Fig. 5f, left). Similar to the free aggression, we observe that the kernel of the VMHvl shell^vgat+^ population increased timed to the onset of attack and the entrance of the mouse does not significantly contribute to the signal change. We observed that the MPO-VMHvl^vgat+^ population is activated by the exit of the mouse (relative to the entrance), and by the offset of attack (Fig. 5f; t(4) = 4.0879, p = 0.015, n = 5 for exit v.s. entrance; Tukey-Krammers post hoc test, corrected p = 0.048 for before attack onset v.s. after attack offset), suggesting that when these events are coincident, they will maximally increase the signal.

**Figure 5.**
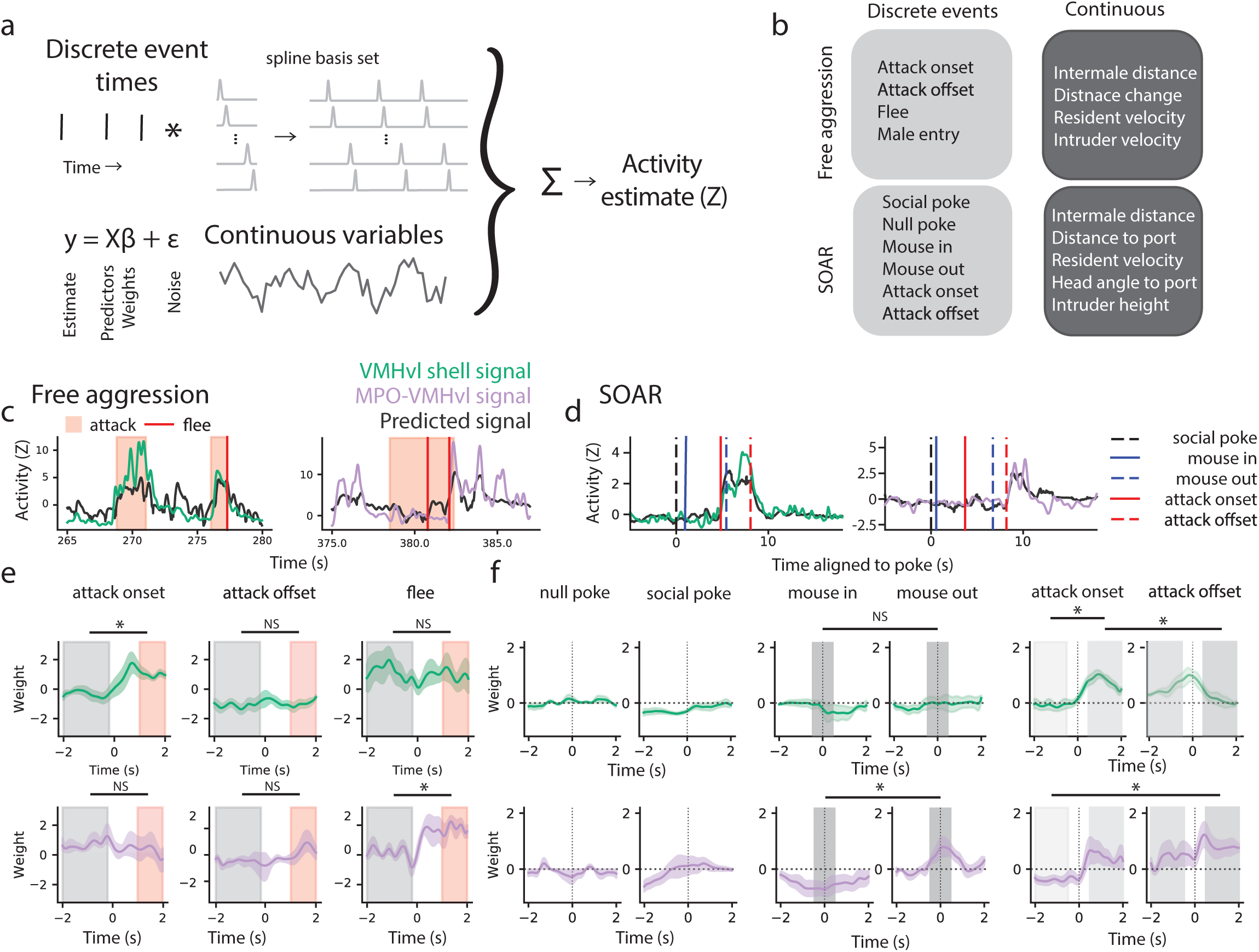
Encoding model reveals that local and long-range inhibitory inputs to the VMHvl are activated by aggressive-action and aggression-ending events respectively. **a-b,** Schematic of the linear encoding model to predict the GCaMP6f signals from Fig. 4. Inputs to the model are the discrete events (**b,** left) convolved with a spline basis set and the continuous features derived from pose tracking (**b,** right). **c-d,** Example of the observed and predicted (gray) GCaMP6f signal during a free aggression for VMHvl shell^vgat+^ (green) and MPO-VMHvl^vgat+^ (purple) for free aggression (**c**) and during the SOAR task (**d**). **e,** Kernel weights for attack onset, attack offset, and flee behaviors during free aggression. The baseline window (gray) is −2 to 0 sec and the post window (red) is 1 to 2 sec from each event. The AUC for the post window was significantly higher than for the baseline window for the attack onset kernel for the signal from the VMHvl shell^vgat+^(t(5) = −3.83, p = 0.012) but not for MPO-VMHvl^vgat+^ (t(4) = 0.514, p = 0.642). The AUC for the post window was not significantly different from the baseline window for the attack offset kernel for either VMHvl shell^vgat+^ (t(5) = − 0.441, p = 0.668) or MPO-VMHvl^vgat+^ (t(4) = −2.165, p = 0.119). The AUC for the post window was significantly higher than for the baseline window for the intruder’s fleeing kernel for the signal from the MPO-VMHvl^vgat+^ (t(4) = −3.395, p = 0.0426) but not for the VMHvl shell^vgat+^ (t(5) = 0.293, p = 0.781). **f,** Kernel weights for discrete events in the SOAR task. The AUC for the ‘mouse in’ kernels was higher than for the ‘mouse out’ kernels for MPO-VMHvl^vgat+^ signal but not for VMHvl shell^vgat+^ (VMHvl shell^vgat+^: t(5) = 0.862, p = 0.428; MPO-VMHvl^vgat+^ : t(4) = 4.0879, p = 0.015). Kernels for attack onset and attack onset show different windows of activation (right). Repeated-measures ANOVA on pre and post attack onset and offset (VMHvl shell^vgat+^ : F_3, 15_ = 6.502, p = 0.0049; MPO-VMHvl^vgat+^ : F_3, 12_ = 4.314, p = 0.0278. VMHvl shell^vgat+^ Tukey-Kramer post hoc test, corrected p = 0.0059, attack onset pre window versus attack onset post window; p=0.0431, attack onset post window versus attack offset post window. MPO-VMHvl^vgat+^ Tukey-Kramer post hoc test, corrected p = 0.048, attack onset pre versus attack offset post). For all panels *p<0.05, **p<0.01, ***p<0.001. Plots show mean±SEM.

Finally, we also tested whether the weights for the continuous variables were significantly different from zero and found that only the weight for the resident velocity during the SOAR task for VMHvl shell^vgat+^ was significantly positive, suggesting that most of the continuous variables did not account for signal change beyond what is accounted for by our discrete events and their time-varying kernels (free aggression: p = 0.0611 for all the four continuous variables for the VMHvl shell^vgat+^, N = 6; p = 0.884, 0.936, 0.694, 0.884 for the intermale distance, distance change, resident velocity, intruder velocity, respectively, for MPO-VMHvl^vgat+^, N = 5; one-sample t-test with Holm-Sidak multicomparison correction) (SOAR task: p = 0.786, 0.798, 0.786, 0.0249, 0.798 for the intruder height, distance to the port, head angle to port, resident velocity, intermale distance, respectively, for VMHvl shell^vgat+^, N = 6; p = 0.996, 0.995, 0.993, 0.996, 0.239, 0.996 for MPO-VMHvl^vgat+^). Overall, these data support an action-related role for the VMHvl shell^vgat+^ population and a motivation-related role for the MPO-VMHvl^vgat+^ population, since activity is maximally driven by behavioral events that signal a break from attack itself.

### Closed-loop optogenetic activation of local and long-range inhibitory populations produces distinct effects on aggressive motivation and action

Since VMHvl neurons encode the temporal sequence of aggression motivation to action, and local and inhibitory neurons that provide inputs to these regions have activity patterns that are active during different phases of this sequence, we next tested whether these phases of aggression could be separately controlled. We injected separate cohorts of vgat-ires-cre males with AAV5-Ef1a-DIO-hChR2(H134R)-mCherry either in the VMHvl shell with a fiber implanted over the VMHvl core (Fig. 6a, Supplementary Fig. 4a) or in the MPO with a fiber over the VMHvl (Fig. 6b, Supplementary Fig. 4b). Control animals were injected with AAV2-Ef1a-DIO-mCherry. All males used in this experiment were sexually and parentally experienced.

**Figure 6.**
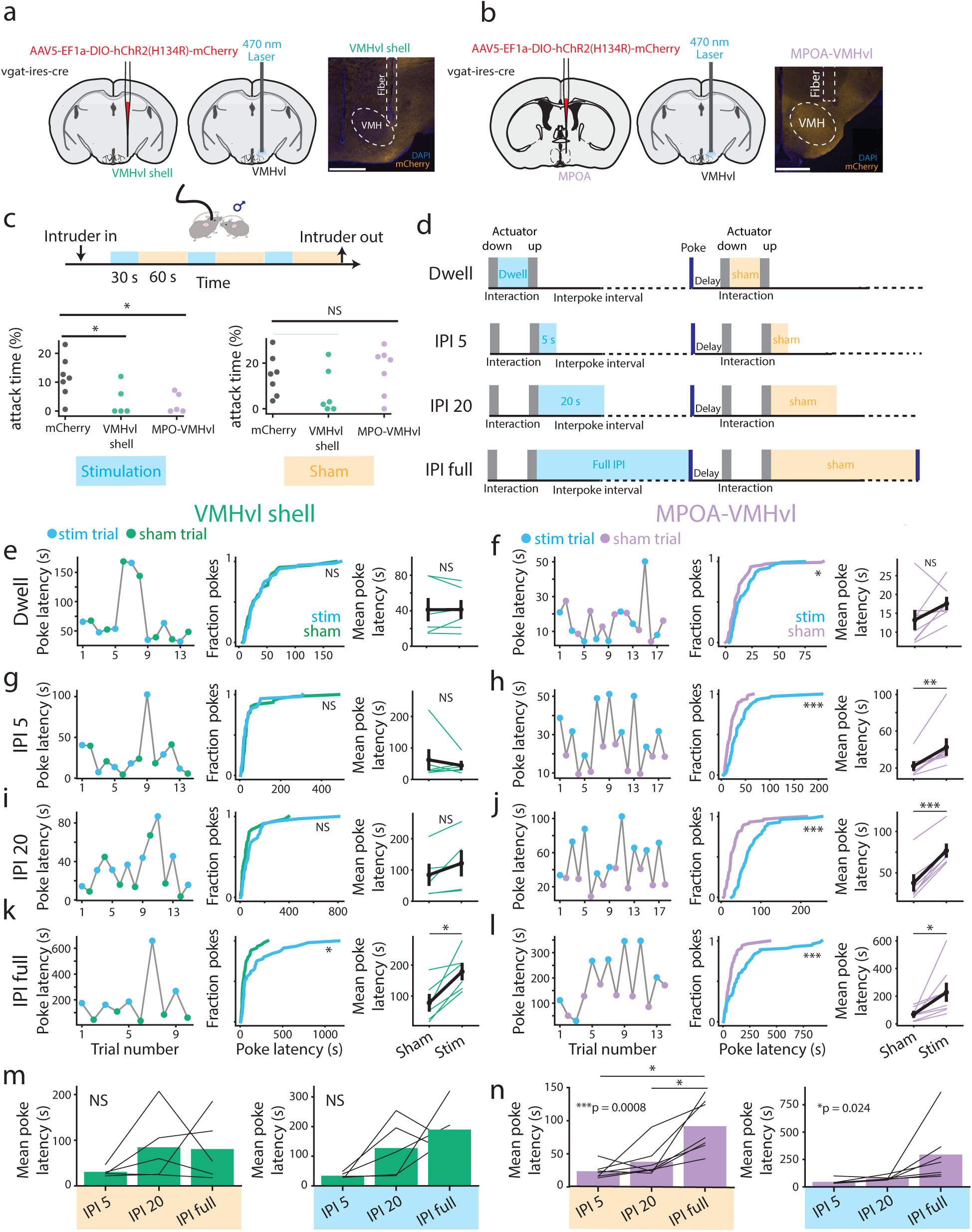
Long-range and local inhibitory inputs to the VMHvl differentially control aggressive motivation and action. **a,b,** Viral strategy for optogenetic activation of VMHvl shell^vgat+^ (**a**) and MPO-VMHvl^vgat+^ (**b**) neurons and histological images showing the fiber location. Scale bars (white), 500 µm. Fiber tips, 200 µm diameter. **c,** Optogenetic activation during free aggression. Percent time in attack was decreased for VMHvl shell^vgat+^ and MPO-VMHvl^vgat+^ relative to no-opsin control animals during the stimulation epochs (F = 6.762, p =0.0069, one-way ANOVA; t(11) = 2.482, p = 0.0305 for control vs shell; t(12) = 3.288, p=0.0129 for control vs MPO-VMHvl, unpaired t-test with post-hoc Holm-Sidak multi comparison correction), but not during the no-stimulation epochs (F =1.517, p = 0.247, one-way ANOVA). N = 7, 6, 7 for no-opsin control, shell, MPO stimulation, respectively. **d,** Closed-loop stimulation schedules during the SOAR task to target the action phase (dwell) and the motivation phase (IPI 5, IPI 20, IPI full). Dwell stimulation lasts from male in to male out (actuator start), and IPI 5, 20, and full stimulation schedules are all triggered on male out (actuator stop) and last for 5s, 20s, or until the time of the next poke, respectively. **e-l,** The effects of optogenetic stimulation during the SOAR task on the poke latencies for VMHvl shell^vgat+^ (**e,g,i,k**) and MPO-VMHvl^vgat+^ (**f,h,j,l**). The left panels show the poke latencies in representative trials for each stimulation schedule. The blue dots represent the poke latencies for stimulation trials and the green (VMHvl shell^vgat+^ or purple (MPO-VMHvl^vgat+^) dots represent the poke latencies for sham trials. The center panels show the cumulative fractions of poke latencies across all the shell or MPO stimulated animals. The right panels show the poke latencies averaged across trials within the session. Each thin green or purple line represents each animal and the solid black line represents the mean across animals with the vertical line showing the standard errors. **e,** Poke latencies were not significantly different between the stimulation trials and the sham trials for the “dwell” regimen for the shell-stimulated animals. Per trial comparison (middle): N = 52 for stim trials and 49 sham trials, p = 0.834, KS statistics = 0.115, Kolmogorov–Smirnov test. Per animal comparison (right): N = 7, t(6) = 0.828, p = 0.439, paired t-test. **f,** Poke latencies were longer for the stimulation trials than for the sham trials in a trial-wise comparison. Per trial comparison (middle): N t= 94 for stim trials and 91 sham trials, p = 0.017, KS statistics = 0.222, Kolmogorov–Smirnov test. Per animal comparison (right): N = 8, t(7) = − 1.956, p = 0.091, paired t-test. **g,**Poke latencies were not significantly different between the stimulation trials and the sham trials for the “IPI 5” schedule of shell-stimulated animals. Per trial comparison (middle): N = 45 for stim trials and 45 sham trials, p = 0.480, KS statistics = 0.178, Kolmogorov–Smirnov test. Per animal comparison (right): N = 7, t(6) = 1.132, p = 0.301, paired t-test; **h,** Poke latencies were significantly longer for the stim trials than for the sham trials for the “IPI 5” schedule of MPO-stimulated animals. Per trial comparison (middle): N = 70 for stim trials and 68 sham trials, p = 0.000032, KS statistics = 0.391, Kolmogorov–Smirnov test. Per animal comparison (right): N = 8, t(7) = −3.814, p = 0.007, paired t-test. **i,** Poke latencies were not significantly different between the stimulation trials and the sham trials for the “IPI 20” schedule of shell-stimulated animals. Per trial comparison (middle): N = 43 for stim trials and 39 sham trials, p = 0.073, KS statistics = 0.274, Kolmogorov–Smirnov test.Per animal comparison (right): N = 6, t(5) = −2.590, p = 0.050, paired t-test. **j,** Poke latencies were significantly longer for the stim trials than for the sham trials for the “IPI 20” schedule of MPO-stimulated animals. Per trial comparison (middle): N = 57 for stim trials and 59 sham trials,t p = 1.69 × 10^-12^, KS statistics = 0.657, Kolmogorov–Smirnov test. Per animal comparison: N = 7, t(6) = −9.783, p = 0.0000656, paired t-test. **k,** Poke latencies were significantly longer for the stim trials than for the sham trials for the “IPI full” schedule of shell-stimulated animals. Per trial comparison (middle): N = 49 for stim trials and 48 sham trials, p = 0.028, KS statistics = 0.282, Kolmogorov–Smirnov test. Per animal comparison: N = 7, t(6) = −4.171, p = 0.006, paired t-test. **l,** Poke latencies were significantly longer for the stim trials than for the sham trials for the “IPI full” schedule of MPO-stimulated animals. Per trial comparison (middle): N = 58 for stim trials and 57 sham trials, p = 0.0000466, KS statistics = 0.413, Kolmogorov–Smirnov test. Per animal comparison (right): N = 8, t(7) = −2.651, p = 0.033, paired t-test; **m,n,** Comparing the mean poke latencies for sham and stim trials across “IPI 5”, “IPI 20”, and “IPI full” stimulation schedules. **m,** (left) Mean poke latencies were not significantly different between the stimulation schedules for the sham trials from shell-stimulated animals (F_2, 8_ = 1.191, p = 0.353, repeated-measures ANOVA, right) The mean poke latencies were not significantly different between the stimulation schedules for the stim trials from shell-stimulated animals (F_2, 8_ = 4.376, p = 0.520, repeated-measures ANOVA, left). **n,** (left) The mean poke latencies were significantly different between the stimulation schedules for the sham trials from MPO-stimulated animals (F_2, 12_ = 13.81, p = 0.0008, repeated-measures ANOVA). By post-hoc test, The mean poke latency for the sham trials was significantly longer for “IPI full” than for “IPI 5” and for “IPI 20”. (t(7) = −4.376, corrected p = 0.012 for “IPI full” v.s. “IPI 5”; t(7) = −3.895, corrected p = 0.012for “IPI full” v.s. “IPI 20”). (right) The mean poke latencies were significantly different between the stimulation schedules for the stim trials from shell-stimulated animals (F_2, 8_ = 5.173, p = 0.024, repeated-measures ANOVA). For all panels *p<0.05, **p<0.01, ***p<0.001.

We first tested the effects of the stimulation of the inhibitory VMHvl shell^vgat+^ and MPO-VMHvl^vgat+^ inputs on aggression during free aggression (Fig. 6c). We delivered stimulation in three 30s bouts (spaced by 60s) during the 5 minute tests. We waited for 30s before delivering the first bout to make sure that the animals had already contacted the intruder before the stimulation. We compared the percentage of time spent in attacking during the three stimulation bouts (90 s in total) between the mCherry control, shell, and MPO-VMHvl stimulated animals, and found that the time spent attacking was significantly shorter for both the shell and MPO-VMHvl group than for the control group (Fig. 6c bottom left; F = 6.762, p =0.0069, one-way ANOVA; p = 0.0305 for control vs shell; p=0.0129 for control vs MPO-VMHvl; n = 7, 6, 7 for no-opsin control, shell, MPO-VMHvl respectively, post-hoc Holm-Sidak multi comparison test). In contrast, the percentage of time spent attacking during the three non-stimulation bouts was not significantly different across groups (Fig. 6c bottom left; F =1.517, p = 0.247, one-way ANOVA). These findings indicate that the stimulation of both the VMHvl shell and the MPO-VMHvl are capable of decreasing the total amount of aggressive behavior.

While we find that stimulating both local and long range inhibitory inputs similarly decreases aggression during free aggression, it is still unclear whether this decrease occurs as a result of decreased aggressive motivation or action, since these processes are not clearly delineated during free aggression. To test more specifically for effects on motivation and action, we conducted closed-loop stimulation during the SOAR task, timing stimulation of local and long range inhibitory inputs to different phases of the self-paced task (Fig. 6d). To test for effects on aggressive action, stimulation was timed to the interaction (“dwell”) phase of the task, since the actions for aggressive motivation had already been complete. To test for effects on aggressive motivation, we performed stimulation during the interpoke interval (IPI) timed to the automated exit of the social target. We used a series of increasing stimulation durations that lasted either 5s (IPI 5), 20s (IPI 20), or lasted the duration of the IPI and ended only when the experimental animal poked the social port again (IPI full). For all stimulation regimes, stimulation was interleaved with sham stimulation on alternating trials.

Previous data demonstrated that activating the VMHvl core during the IPI increased aggressive motivation and decreased poke latency on stimulation trials relative to sham stimulation trials^8^. To test whether inhibitory inputs have a role in determining aggressive motivation, we reasoned that activation of these inputs would increase the latency to the next poke (Fig. 6e-l). For each stimulation regime we compared the full distribution of poke latencies for stimulation and sham trials, and also the mean latency for stimulation and sham trial for each animal. While we observed that stimulation during the dwell period did not significantly change the poke latency for either local or long-range inhibitory inputs (Fig. 6e-f; VMHvl shell: t(6) = 0.828, p = 0.439, n = 7; MPO-VMHvl: t(7) = −1.956, p = 0.091, n = 8; paired t-test), we observe robustly different patterns of behavior for stimulation regimes during the IPI. Surprisingly, we observe that for both IPI 5 and IPI 20, poke latencies following stimulation trials are significantly longer for MPO-VMHvl^vgat+^ (t(7) = −3.814, p = 0.007, n = 8 for IPI 5; t(6) = −9.783, p = 0.000066, n = 7 for IPI 20; paired t-test) but not for the VMHvl shell^vgat+^ stimulation (t(6) = 1.132, p = 0.301, n = 7 for IPI 5; t(5) = −2.590, p = 0.050, n = 6 for IPI 20; paired t-test) (Fig. 6 g-j, for each panel see example session, left, comparison across trials, middle, and comparison across animals). This demonstrates that even brief stimulation, when timed to the end of the aggressive interaction in MPO-VMHvl^vgat+^ can generate long-lasting behavior effects. Only when stimulation lasted all the way to the next poke (IPI full), did we observe effects on aggressive motivation with both VMHvl shell^vgat+^ and MPO-VMHvl^vgat+^ stimulation (Fig. 6k-l; VMHvl shell: t(6) = −4.171, p = 0.006, n = 7; MPO-VMHvl: t(7) = −2.651, p = 0.033, n = 8; paired t-test). No stimulation regimes significantly affected the latency to the next poke in control animals (Supplementary Fig. 4c-f; paired t-test, p = 0.141 dwell; p=0.32, IPI 5; p=0.492 IPI 20; p=0.487 IPI full; n = 7; paired t-test). We also observed that increased duration of the stimulation also increased the poke latencies of sham stimulation trials interleaved with the real stimulation trials for the MPO-VMHvl^vgat+^ but not VMHvl shell^vgat+^ stimulation trials (Fig. 6m-n; VMHvl shell: F_2, 8_ = 1.191, p = 0.353, MPO-VMHvl: F_2, 12_ = 13.81, p = 0.0008; repeated-measures ANOVA). This suggests that stimulation of the MPO-VMHvl^vgat+^ may have uniquely long lasting effects that remain even after the stimulation itself is over, pushing the animals into an unmotivated state.

In contrast, we observed that VMHvl shell^vgat+^ stimulation, but not MPO-VMHvl^vgat+^ stimulation resulted in a change in aggressive action. VMHvl shell^vgat+^ stimulation during the “dwell” phase significantly reduced the amount of time spent attacking during that interaction (Fig. 7a, t(5) =3.627, p = 0.0151, n = 6, paired t-test), while stimulation of MPO-VMHvl^vgat+^ did not significantly change time spent attacking (Fig. 7b, t(5) = 0.864, p = 0.4269, n = 6, paired t-test). Overall, these data demonstrate a double dissociation of function for local and long-range inhibition to the VMHvl: long-range inhibition from the MPO decreases motivation, reflected in increased poke latency and promoted an unmotivated state, but did not affect attack itself. In contrast, stimulation of the local inhibitory input did not change aggressive motivation, except when stimulation lasted all the way until the poke itself, but significantly decreased attack action.

**Figure 7.**
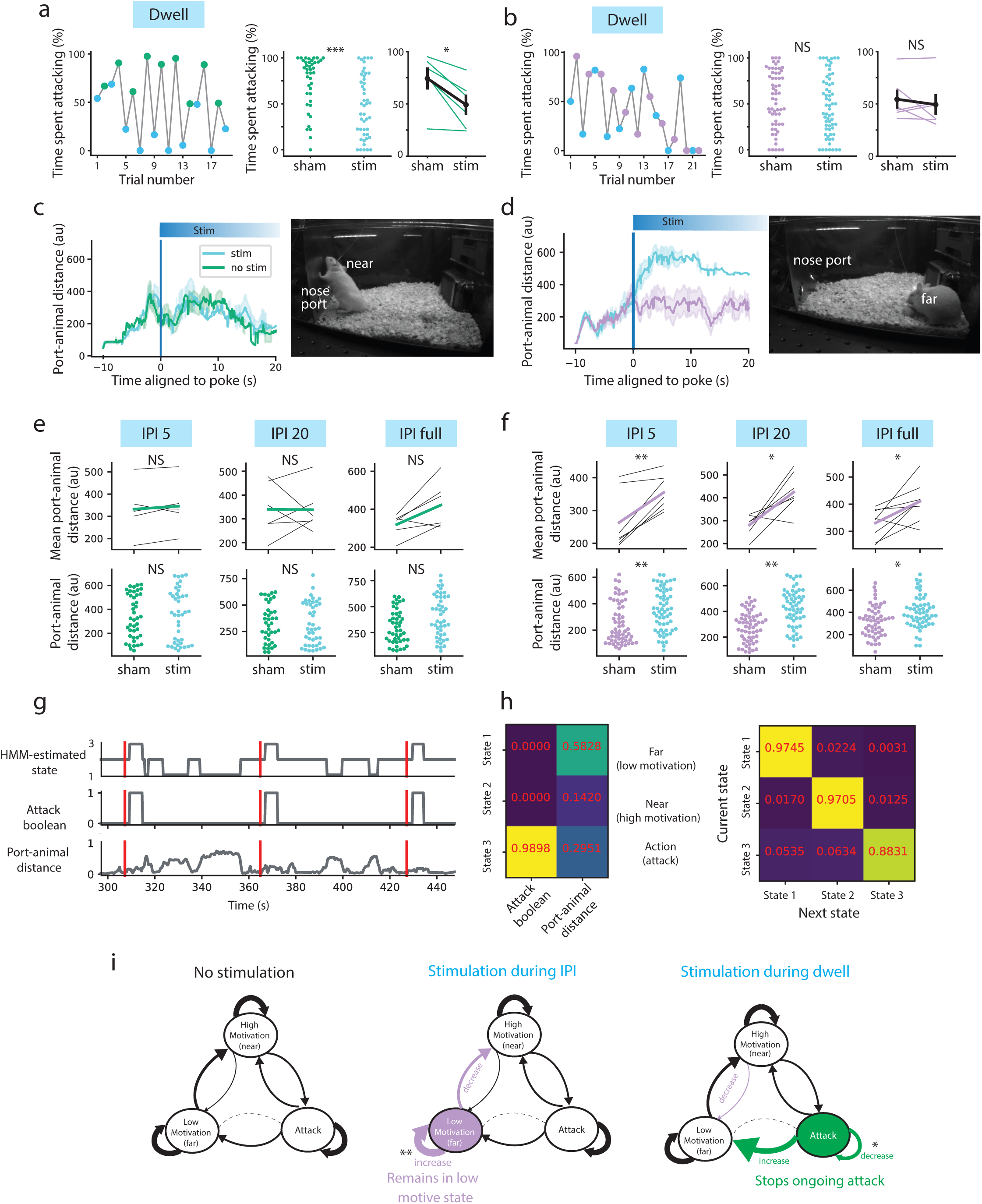
Long range and local inhibition differentially modify instantaneous behavior and change aggressive state. **a, b,** The effects of optogenetic stimulation for the “dwell” schedule on the time spent attacking. Left, the percentage of time spent attacking in a representative session. Center, the distribution of the percentage of time spent attacking through all the animals. Right, the percentage of time spent attacking averaged within a session for each animal. **a,** The “dwell” stimulation of the VMHvl shell^vgat+^ significantly decreased the time spent attacking. The distribution of the percentage of time spent attacking for stim was significantly different from that of sham trials (p = 0.0004, N = 46 stim trials and 41 sham trials, Kolmogorov-Smirnov test). The averaged percentage of time spent attacking per animal was significantly smaller for the stim trials than for the sham trials (t(5) =3.627, p = 0.0151, n = 6, paired t-test). **b,** The “dwell” stimulation of the MPO-VMHvl^vgat+^ did not significantly change the time spent attacking. The distributions of the percentage of time spent attacking for stim versus sham trials were not significantly different (p = 0.3165, N = 63 stim trials and 59 sham trials, Kolmogorov-Smirnov test). The averaged percentage of time spent attacking per animal was not significantly different between the stim trials and the sham trials (t(5) = 0.864, p = 0.4269, n = 6, paired t-test). **c, d,** The effect of optogenetic stimulation during the interpoke interval on the animal’s nose-to-port distance. Distance change under “IPI full” stimulation schedule. Right, a representative photo of a side camera view in which the experimental animal is located (**c**) near to the nose port or (**d**) far from the nose port on the left corner of the cage. **e,f,** Mean nose-port distance for the first 20 seconds of the interpoke interval. Top row shows the mean distance per animal and the bottom row shows the mean distance per trial. **e,** The distance to the social port averaged for the first 20 seconds was not significantly different between the stim and the sham trials for all the stimulation schedules for VMHvl shell^vgat+^ stimulation. Per animal comparison: t(5) = −1.088, p = 0.326 for IPI 5, t(5) = 0.0369, p = 0.972 for IPI 20, t(5) = −2.538, p = 0.052 for IPI full, N = 6. Per trial comparison: p = 0.667, N = 41 stim trials and 42 sham trials, KS statistics = 0.151 for IPI 5; p = 0.792, N = 43 stim trials and 39 sham trials, KS statistics = 0.134 for IPI 20; p = 0.081, N = 45 stim trials and 45 sham trials, KS statistics = 0.267 for IPI full. Paired t-test for per animal comparison and Kolmogorov-Smirnov test for per trial comparison. **f,** The distance to the social port averaged for the first 20 seconds was significantly farther for the stim trials than for the sham trials for all the stimulation schedules for MPO-VMHvl^vgat+^ stimulation. Per animal comparison: t(6) = −4.492, p = 0.004 for IPI 5, t(6) = −3.575, p = 0.0117 for IPI 20, t(7) = 2.853, p = 0.025 for IPI full. N = 7 for IPI 5 and IPI 20 and 8 for IPI full. Per trial comparison: p = 0.003, N = 60 stim trials and 59 sham trials, KS statistics = 0.325 for IPI 5; p = 2.99 × 10^-6^, N = 57 stim trials and 59 sham trials, KS statistics = 0.466 for IPI 20; p = 0.0143, N = 56 stim trials and 55 sham trials, KS statistics = 0.293 for IPI full. Paired t-test for per animal comparison and Kolmogorov-Smirnov test for per trial comparison. **g**, Example 2-feature, 3-state HMM model of the stimulation dataset **h,** Emission matrix (left) of the model and transition matrix (right) of the 3-state HMM model. **i,** Schematic summarizing the results of comparing the different transitions between the stimulation and sham-stimulation data. For all panels *p<0.05, **p<0.01, ***p<0.001. Plots show mean±SEM.

We also used behavioral quantification to test whether we could observe behavioral correlates of an unmotivated state during IPI stimulation. Specifically, we reasoned that if stimulation generated an unmotivated state, mice would spend less time close to the “high-motivation” cage location where they request more aggression. We quantified the distance from the animal’s head to the nosepoke port and found that VMHvl shell^vgat+^ stimulation did not change the likelihood of the animal changing his location on a trial-to-trial basis (Fig. 7c,e; t(5) = −1.088, p = 0.326 for IPI 5, t(5) = 0.0369, p = 0.972 for IPI 20, t(5) = −2.538, p = 0.052 for IPI full, N = 6). In contrast, we observed a significant shift in this distance with MPO-VMHvl^vgat+^ stimulation (Fig. 7d,f, t(6) = −4.492, p = 0.004 for IPI 5, t(6) = −3.575, p = 0.0117 for IPI 20, t(7) = 2.853, p = 0.025 for IPI full, N = 7 for IPI 5 and IPI 20 and 8 for IPI full), suggesting that stimulation evokes instantaneous behaviors that reflect an unmotivated state.

Finally, to further illustrate this difference between an unmotivated and motivated state, we fit a Hidden Markov Model (HMM) to this data during stimulation and sham stimulation (Fig. 7g-h)^27,28^. We fit a series of models with 2-5 states model to a two feature model consisting of the continuous distance between the animal and nosepoke port, and a Boolean vector representing whether that time point contained attack, and found using BIC that the increasing beyond 3 states did not significantly improve the model (using BIC, Supplementary Fig. 5a). The 3-states of the model (as defined by the resulting emission matrix, Fig, 7h, left) represented states where the animal was far and not attacking (state 1: low motivation state), close and not attacking (state 2: high motivation state), and attacking (state 3: action state). We next calculated the transition probabilities for the full dataset (Fig. 7g) and separately for each stimulation regime and compared the change in transition between each of the states (Supplementary Fig. 5b-c). We found that in trials with no stimulation (schematized in Fig. 7i, left), transitions within-state were the most probable, but animals occasionally transitioned between states (though seldom transitioned directly from a low motivation state directly to attack, Fig. 7h, right, Fig. 7i left, dotted line). We compared these transition probabilities to updated probabilities for stimulation during the action phase (dwell) and during the motivational phase (IPI 5 and IPI 20). We found that for stimulation during the motivational phase, MPO-VMHvl^vgat+^ stimulation significantly increased the probability that the animal remained in the low motivational state and decreased the likelihood of transitioning into the high motivated state (Fig. 7i, middle). VMHvl shell^vgat+^ stimulation does not significantly change transition likelihoods (Supplementary Fig, 5b-c). In contrast, for stimulation during the dwell phase, VMHvl shelll^vgat+^ stimulation increases the likelihood of transitioning out of attack (Fig. 7i, right). These data illustrate the distinct effects of these two gating mechanisms: Activation of a long-range inhibitory input can prolong a distinctly low motivational state, and can delay the transition to a high-motivational state, while activation of a local input can stop ongoing attack, and promote the exit from an action state.

## Discussion

Here, we demonstrate that the VMHvl has a key role in encoding the transition of aggressive motivation to aggressive action, and that these processes can be independently gated through separable inhibitory inputs. We find that neurons in the VMHvl reliably encode motivation-related signals that precede attack, irrespective of the specific actions that comprise the motivational phase. In addition, we find that two key inhibitory inputs (local inhibition from the VMHvl shell and long-range input from the MPO) have distinct temporal patterns of activation, whereby local inhibitory inputs are action-selective, and long-range inhibitory inputs are activated at the offsets of these interactions, in particular, when the aggressive target moves away, signaling the end of an attack event. Finally, we find that local inhibition can specifically gate the action phase of aggression, while long-range inputs decrease motivation by preventing the re-initiation of an aggressive state.

We do not find evidence for a motivation-specific and action-specific “labeled line”, since we find that both local and long-range inputs promiscuously target heterogeneous VMHvl core neurons (Fig 2). While this data suggests that local and long-range neurons do not synapse onto different postsynaptic populations, they do not rule out specificity within these projections (i.e. that subsets of this projection have more specific projection patterns to subsets of VMHvl core neurons). However, in both our *ex vivo* slice physiology and in our optogenetic-identity tagging experiments, we find both direct and indirect evidence for different weights from the MPO (but not the VMHvl shell). This suggests that inhibitory MPO neurons may be strongly coupled neurons in the posterior part of the VMHvl core which contains more motivation-preferring neurons.

Additionally, we find that local and long-range inhibitory inputs to the VMHvl have distinct activity profiles during aggressive interactions, suggesting that their ability to gate or modify behavior may be limited to those timepoints. While these populations display critical differences in the timing of activity relative to attack, the inputs also share two key properties. First, activity profiles at the moment of the poke action are both low, suggesting that they may both play a permissive role in the initiation of the motivated action: the poke. This is confirmed by the fact that activation of the VMHvl shell population, like activation of the MPO-VMHvl projection, up to the moment of the poke can both delay poke initiation. Second, both populations are robustly activated by pup investigation. Though virgin males will frequently attack pups, often resulting in infanticide^29^, sexually experienced, parentally experienced males seldom do^30^. Parental behavior has been shown to induce c-fos in the MPO and lesions of the MPO can convert a parental father to an infanticidal one^30^. Since pup investigation can activate both inputs, we interpret this as a “double locking” mechanism such that gating mechanisms prevent both motivation and action in conditions where attack would be severely inappropriate. The fact that we do not observe decay in the response of the MPO-VMHvl^vgat+^ population across repeated investigation episodes further suggests that this input plays an important role in preventing the transition to a high-motivated state.

Surprisingly, despite extensive previous data showing that the MPO plays a critical and highly conserved role in male mating behavior^16,31^, our data suggests that the MPO-VMHvl^vgat+^ population does not play the same strong “double-locking” role in attack suppression during investigation of a female as we observe during pup investigation. (However, activity is very modestly increased in the MPO-VMHvl population timed to the female’s entrance.) However, since the VMHvl receives inhibitory input from a variety of other regions known to contain sex-biased coding^5,32^, these other regions may perform other specific types of context gating to prevent attack in the presence of a female.

We observe that neurons in the VMHvl are activated in a sequence from motivation to action during aggression and that this sequence does not depend on the actions in the motivational phase. We do not think that this sequence emerges locally, as a result of a synfire chain activated through local feed-forward excitation, since rates of local synaptic connectivity in adulthood are extremely sparse^14^ (although there may be extensive recurrent activity with neighboring hypothalamic partner the PMv^33^). Instead, this sequence may instead emerge as a consequence of specific spatiotemporal windows of inhibition and disinhibition^34^. In our data, both inputs behave permissively during the motivational phase (the poke), while acute local inhibition is able to titrate the duration of attack. Our data suggests that brief inhibition timed to the offset of attack (but not during attack itself) can have long lasting suppressive effects on aggressive motivation, since individuals enter an acutely unmotivated state and do not perform aggression-seeking actions. Since both the MPO and VMHvl are highly molecularly heterogeneous^24,35^, one possibility is that this long-lasting effect, in some cases, many minutes, may be mediated through peptidergic transmission. Recent modeling work has suggested that peptide-like signaling is required for the persistence of neural and behavioral phenotypes in predator fear circuits^36^. Such long-lasting mechanisms might be evolutionarily advantageous as they allow the prioritization of other behaviors over aggression in a computationally inexpensive manner.

The inhibitory input from the MPO to the VMHvl has a simple relationship with behavior output: when activity is low, it behaves permissively allowing the initiation of an aggressively motivated state, and when activity is high, it prevents this initiation, with a long decay. However, the VMHvl shell has a more complex relationship with its postsynaptic partner, the VMHvl core. Previous data has implicated the VMHvl shell to VMHvl core circuit in the circadian regulation of aggression^18^, but, unlike the MPO, the activity of the core and shell do not have an inverse relationship. Instead, our optotagging and photometry recording experiments reveal that activation peaks are later than neurons in the posterior VMHvl, suggesting that the activity of these neurons may behave as a deceleration mechanism once the motivation-action sequence is already in progress. Inhibition of the downstream attack actuator mechanisms in the lPAG have similar action-specific effects^37^.

One major open question is how flexible these circuit functions are with age and experience. Here, we capture these neural dynamics and putative synaptic weights in the highly trained, aggressively, sexually, and parentally experienced animal. Given the flexibility of these circuits, it is likely that these properties change as a function of experience, hormones, and experience-dependent hormones^38,39^. Experience-dependent flexibility can manifest as change in the inputs, the strength of the connectivity, or in changes to understudied differences in hormone levels and their receptors. Use of hidden Markov models or other latent state models that allow motivational and other internal states to fluctuate on multiple timescales will be critical to capturing this long-term variability^40^.

Overall these data demonstrate that distinct circuit-level control mechanisms on aggressive motivation and action can modify instantaneous and future behavior in separable ways. While many studies of social behavior describe behavioral change along a single axis (e.g. aggression increases or decreases), here we show that the brain actually contains several types of braking mechanisms that adjust circuit functionality in different ways. Together, these data illustrate the power of using more constrained behavioral tasks in tandem with free behavioral assays to better segregate behavioral difference and circuit function.

## Methods

### Mice

All animal procedures were approved by the Princeton University Institutional Animal Care and Use Committee and were in accordance with National Institutes of Health standards. All experimental animals were male, and sex was assigned based on observation of genitalia at weaning. In fiber photometry recording studies, we used 15 Vgat::Cre with Swiss-Webster background. For *in vivo* optogenetics, we used 16 Vgat::Cre mice and 6 wild type mice. For whole-cell patch clamp experiments, we used 15 Vgat::Cre mice and 1 wild type mouse with Swiss-Webster background. All mice were singly-housed prior to *ex vivo* experiments. Male and female conspecifics used as social targets in freely moving assays and the social operant task were sexually naive, BALB/c (male) or Swiss-Webster (female) mice between the ages of 10 and 36 weeks. Mice were housed in a 12 h light-dark cycle with experiments taking place exclusively during the dark phase. Food and water were given ad libitum. Prior to behavioral testing, the experimental mice were paired with female Swiss-Webster wild type mice 2-5 days after surgeries, and they were kept as breeding pairs until all the experiments were completed. All animals were bred within the Princeton University animal facilities or purchased from Taconic Biosciences.

### Behavior Tests

Animal behavior video was recorded using up to five cameras (BFS-U3-13Y3m-C 1.3 Mono, FLIR Systems, Inc.) acquired with SpinView, and controlled with an RP2.1 or RX8 (Tucker-Davis Technologies, Inc.).

### SOAR task

The nosepoke apparatus for the social operant task with actuator-mediated reward (SOAR) consisted of two custom-made nosepoke ports and a motorized actuator (12-inch Classic Rod Linear Actuator, Firgelli) holding a reward animal controlled by a minicomputer (Raspberry Pi). Each nosepoke port contained a pair of an infrared detector and an emitter on the sides and a yellow LED at the center so that the system could detect a nosepoke and send a visual cue to the animal when an experimental animal nosepoked by turning off the yellow LED transiently. Prior to each training session, nesting materials, food pellets, and other mice (including a female partner and pups) were removed from the home cage. For each session, a metal panel attached to the two nosepoke ports was inserted vertically to the cage. The reward animal, a male, group-housed BALB/c was vertically suspended from a horizontal arm attached to the tip of the actuator. When the experimental animal nosepoked into the right-hand port (relative to the animal when it faced the port), it activated retraction of the actuator to deliver the reward animal into the cage of the experimental animal after a random delay (drawn from a distribution 0 -1s), and after a randomly timed social interaction (0.2-3s), the reward animal exited the cage as the actuator extended. Once triggered, the system could not be triggered again until the reward animal moved back to the original position (“lock-out”). The actuator traveling time was fixed to 4s for both directions. If the experimental animal nosepoked the left-hand ‘null’ port, the system was not activated although the yellow LED in the null port turned off. All the timestamps of nosepokes and actuator movements were recorded by the Raspberry Pi, and also by the RX8 data acquisition board.

Mice were trained for 5-12 successive sessions and animals were considered ‘learned’ if they reached poke rates of at least 0.4 pokes per min for the social port and reached performance levels of greater than 66.6% preference to the social port on at least two successive days.

### Resident intruder

For in vivo physiology recordings, intruder mice (either a submissive BALB/cJ male or a sexually mature SWR/J female, stocks 000651 and 000689) were introduced to the cage in a block format, in pairs of 60 s trials separated by 120 s. To enable data synchronization, a manual button tied to both an in-frame LED and the data acquisition system was pressed as the stimulus animal was introduced to the home cage, and again after 60 s during removal for each trial. For fiber photometry recording, we introduced an intruder animal two minutes after the onset of the calcium recording to allow the recording of baseline calcium activities before the entry of an intruder. Typical resident intruder assays lasted for four minutes, and then the intruder was removed and the recording lasted for another two minutes with the experimental animal staying alone to see any possible signal changes after the resident-intruder assay. The intruder mice were either a submissive BALB/cJ male, a Swiss-Webster female, or a novel Swiss-Webster pup between 1-10 days old from different breeding cages. For the resident-intruder assays for *in vivo* optogenetic stimulation, we introduced an intruder animal (a BALB/cJ male) into the cage and the animals received three bouts of 30-sec stimulation separated by 60-sec intervals, with the first bout started 30 seconds after the intruder entered the stimulated animal’s home cage for a total of 5 minutes.

### Behavioral scoring and classification

Video recordings of both the SOAR behavior and resident-intruder trials were manually scored using Behavioral Observation Research Interactive Software (BORIS v8.8 https://besjournals.onlinelibrary.wiley.com/doi/full/10.1111/2041-210X.12584). Behavioral point events included attacking, grooming, eating, close sniffing, and intruder entry/exit, while state events included bites and batting for the *in vivo* electrophysiology experiments.

For fiber photometry and optogenetic experiments, the point events included intruder entry/exit (resident-intruder), the onset of the session (the SOAR task) and intruder fleeing (resident-intruder). The state events included attacking, close sniffing (referred to as “investigation” in the texts and figures), and eating.

Animal pose estimation was performed with SLEAP v1.1.5 or with DeepLabCut v2.0.6. To create a model, mice in a subset of frames from behavior videos were hand-labeled with a skeleton of 15 nodes. These frames (>6,000) were then used as a training and validation set to create a model (see below). We then used this model to run inference on all behavior videos, whereby the model produced predictions that we then hand-corrected to produce final pose estimation. Data were then analyzed in MATLAB or Python using custom scripts. In a subset of behavior, we used supervised classification to apply behavior behavioral labels (see below).

### *In vivo* electrophysiology during aggression

#### Surgical implantation and recording with the shuttleDrive

Male mice bred from SWR/J and Vgat-ires-cre knock-in C57BL/6J (The Jackson Laboratory, stocks 000689 and 028862) were injected with a viral strategy including 250-400 nL of rAAV2/Ef1a-DIO-eNPHR3.0-EYFP (UNC GTC Vector Core, titer of 4.1×10^12) to MPO (−3.1 A/P, +/-0.5 M/L, −4.8 D/V), and AAV1 virus produced from pAAV-EF1a-double floxed-hChR2(H134R)-mCherry-WPRE-HGHpA (addgene 20297, titer of 1.8×10^13) to VMHvl (−4.65 A/P, +/-0.75 M/L, −5.8 D/V). Coordinates are listed in mm to Anterior/Posterior, Medial/Lateral, and Dorsal/Ventral, relative to the intersection of the sagittal suture with the inferior cerebral vein, and to brain surface. A polished 200um core fiber (MFC_200/245-0.53_32mm_MF1.25_FLT, Doric Lenses Inc.) was then implanted at a 45 degree lateral angle 0.1 mm above the MPO viral injection. Finally, a craniotomy was opened over VMH and a ShuttleDrive (OpenEphys) loaded with twisted tetrodes (Stablohm 650, California Fine Wire Company) and optical fibers (MFC_200/245-0.53_32mm_MF1.25_MA60, Doric Lenses Inc.) was implanted at a depth of −5.6 mm. The animal was allowed to recover, and tetrodes were advanced until laser-responsive units were recorded.

At the end of a behavior session, optogenetic identify-tagging experiments consisted of pulsing 5 ms 2-4 Hz pulses of 5 mW blue light to the VMH, and connectivity with inhibitory inputs was determined using 500 ms 0.1 Hz pulses of 10 mW yellow light to the MPO.

Recordings were digitized with a DigitalLynx SX (Neuralynx) at 30 kHz using an HS-64-MUX headstage and Saturn 2 commutator, and acquired using Cheetah 6.3.2. Blue and Yellow optical stimulation was delivered with an MDL-III-460 and MGL-FN-556, respectively (Opto Engine LLC) via a FRJ_1×2i_FC-2FC rotary joint (Doric Lenses Inc.).

#### Spike sorting, clustering criteria

Spike sorting was performed offline from continuously-recorded channels (band-pass filtered during acquisition at 300-7500Hz) based on relative spike amplitude, energy, kinetics, and principle component space using Plexon Offline Sorter v4 following minor post-processing, which included subtraction of median optical artifacts during blue light stimuli and de-medianing to conservatively remove common movement artifacts. Units were considered well-isolated when they had spike heights more than double the rms noise, were unimodally distributed across feature spaces, and when clusters were well-separated from both each other and the unity line in energy space that occurs due to residual movement artifact crossing threshold.

#### Analysis of neural data

All analyses for spiking data were performed using custom code written in Matlab. For population response of neural activity, spike trains were smoothed with a 25ms gaussian window on each trial, then for plotting purposes, averaged across trials and smoothed with a 300ms sliding window (using 100ms bins) and z-scored. Selectivity indices for motivation and action were computed on raw firing rates with no smoothing. Selectivity indices were computed using the equation Selectivity = (A-B)/(A+B) where A and B represent the mean activity for a specific time bin for each neuron. Motivation selectivity was defined as the mean activity in the window from −1 prior to nosepoke to the actuator down time, relative to a baseline window during the interpoke interval (male out +1s to −8 prior to nosepoke). Action selectivity was defined by mean activity from the actuator down (male in) to actuator up (male out) relative to the baseline. Motivation-action selectivity was defined as the motivation epoch relative to the action epoch. Peak times relative to attack onset were computed on smoothed mean responses across trials. Early and late motivation neurons were defined as having their peak - 1.75 to -1s and -1s to -0.25s prior to attack onset respectively. Onset neurons are −0.25s to 0.25s around to attack onset, and action neurons are 0.25 to 1.75 after attack.

Units were defined as being VMHvl shell “optotagged” or VMHvl suppressed if they were significantly increased or decreased (respectively) by comparing the trial-to-trial response in the 30ms prior to the blue light pulse to the 5ms after the light pulse using a paired t-test and corrected using Benjamini-Hochberg false discovery rate correction. For the “looser” identification criteria (Supplementary Figure 3), we expanded the post-light window to 10ms. Units defined as being “MPO modulated” were significantly increased in the 500ms window following yellow light relative to the 500ms prior to the light pulse (using paired t-test and Benjamini-Hochberg false discovery rate correction). For unsupervised hierarchical clustering of neural responses (Wards), we clustered a matrix of the concatenated z-scored neural responses for each neuron aligned to the introduction of the male or female (−5s to +5s), and to investigation of male or female or attack (−2s to +2s). Only neurons that were recorded in both freely moving behavior with both males and females and also during the SOAR task were used for clustering. We tested for cluster enrichment by identity-tagged neurons by comparing the frequency of the in-cluster neurons to the frequency of the particular class of identified neurons for each cluster using a Fisher’s exact test, corrected for multiple comparisons (5 clusters) using Bonferroni correction.

### Ex vivo whole-cell patch clamp electrophysiology

#### Surgery details

P42 or older male vgat-ires-cre Swiss-Webster background mice (bred as described above) were injected with 200 nl of AAV1-Ef1-DIO-hCHR2(H134R)-eYFP-WPRE-HGHpA bilaterally to MPO (−3.1 A/P, ±0.5 M/L, −4.8 D/V), and 200 nl AAV2-Ef1-DIO-mCherry bilaterally to VMHvl shell (−4.8 A/P, ±0.8 M/L, −5.8 D/V) for experiments involving optical stimulation of MPO terminals and 200 nl of AAV1-Ef1-DIO-hCHR2(H134R)-mCherry-WPRE-HGHpA bilaterally to VMHvl shell for experiments involving stimulation of shell terminals in VMHvl core. The animal was allowed to recover, and experiments were conducted at minimum 5 weeks following surgery.

#### Ex vivo *slice preparation*

Mice were first deeply anesthetized with sodium pentobarbital (Euthasol, Penn Veterinary Supply) and then decapitated following cessation of reflexes. Brains were rapidly dissected and placed in an ice-cold *N*-methyl-D-glucamine (NMDG)-based slicing solution (in mM: 93 NMDG, 2.5 KCl, 1.2 NaH_2_PO_4_, 30 NaHCO_3_, 20 HEPES, 25 Glucose, 5 Na-ascorbate, 2 Thiourea, 3 Na-pyruvate, 12 *N-*Acetyl-L-cysteine [NAC], 10 MgCl_2_, 0.5 CaCl_2_; pH adjusted to 7.3-7.4 using up to 93 HCl). All solutions (slicing, recovery, and ACSF) were kept oxygenated with 95% O_2_-5% CO_2_. 300 μm coronal sections through the entirety of the mediobasal hypothalamic region were obtained using a vibratome (VT1200s, Leica Biosystems, Germany). Following sectioning, slices were first allowed to incubate at 36°C for 12 minutes in the NMDG-based slicing solution before being transferred to a recovery solution (in mM: 92 NaCl, 2.5 KCl, 1.2 NaH_2_PO_4_, 30 NaHCO_3_, 20 HEPES, 25 Glucose, 5 Na-ascorbate, 2 Thiourea, 3 Na-pyruvate, 12 NAC, 2 MgCl_2_, 2 CaCl_2_) for 1 hour at room temperature. All solution reagents were purchased from Sigma Aldrich (St. Louis, MO, USA).

#### Electrophysiology

Slices were placed in a submerged slice chamber and perfused with ACSF (in mM: 124 NaCl, 2.5 KCl, 1.2 NaH_2_PO_4_, 24 NaHCO_3_, 5 HEPES, 12.5 Glucose, 2 MgCl_2_, 2 CaCl_2_) heated to 32-37°C. Slices were visualized with a moving stage microscope equipped with differential interference contrast and fluorescence imaging (Scientifica, Uckfield, UK; Lumencor, Beaverton, OR, USA). Whole cell recordings were made using borosilicate glass pipettes (2.5-7.0 MΩ; Sutter, Novato, CA, USA) filled with intracellular recording solution (in mM: 130 KCl, 5 CaCl_2_, 10 EGTA, 10 HEPES, 2 MgATP, 0.5 Na_2_GTP, 10 Na_2_-Phosphocreatine). To isolate inhibitory currents, 10 μM NBQX and 50 μM D-APV were included in the bath and cells were maintained at V_m_ = −70 mV for current clamp and voltage clamp experiments. Access resistance was monitored throughout experiments and data were discarded if access resistance exceeded 35 MΩ or varied by > 20%. Data were acquired at 10 kHz and lowpass filtered at 4 kHz using a Multiclamp 700B amplifier and were digitized using a Digidata 1440A with pCLAMP 10.7 software (MDS Analytical Technologies, Sunnyvale, CA, USA). Offline, current data were filtered using a 2 kHz lowpass filtered.

#### Cell type identification

VMHvl core neurons were identified as Vgat-mCherry^-^neurons found within the shell of Vgat-mCherry neurons that surround the VMHvl core (Fig. 2a). Position within VMHvl core was visually confirmed post hoc.

#### Experimental design and quantification

##### Optical stimulation experiments

For all optical stimulation experiments, blue light illumination (1-5 ms) was focused to ChR2^+^ axon terminals surrounding the recorded cell using a 40X, 0.8 NA water immersion objective (Olympus, Tokyo, Japan). First, the minimal optical stimulation threshold for each recorded neuron was determined. This was defined as the lowest optical stimulation parameters (combined light intensity and duration) to elicit postsynaptic responses. This minimal stimulation threshold was used for both single stimulation and paired pulse experiments. Recordings were additionally made at the parameters just below the threshold to verify lack of response. Due to the depolarizing nature of chloride with the intracellular solution we used, optical stimulation of Vgat^+^ terminals occasionally led to the initiation of an inward voltage-gated sodium current. Trials in which these currents occurred and obscured the oIPSC were discarded for both single stimulation and paired stimulation experiments.

Single optical stimulation experiments: oIPSCs were recorded in response to a single stimulation of Vgat-ChR2^+^ terminals. 10-30 trials were repeated at 0.067 Hz. oIPSCs were detected using custom MATLAB code with an oIPSC being defined as a negative deflection within 10 milliseconds of optical stimulation that exceeded baseline current by 5X the median absolute deviation of the 500 ms of current data prior to optical stimulation. oIPSC amplitude was computed as the difference between oIPSC peak and the mean of the 500 ms baseline period. The cellular mean oIPSC was computed as the mean of all the successful oIPSCs recorded in the neuron.

Paired pulse experiments: Experiments were conducted the same as the single stimulation experiments except optical stimulation was given in a burst of two pulses of light separated by 20 ms. All trials were then averaged together, and the paired pulse ratio was computed as peak amplitude of the second oIPSC divided by the peak amplitude of the first oIPSC.

##### Current clamp experiments

Square current steps: 1 second long square current injections (−200 pA to +200 pA, Δ20 pA) were given at 0.25 Hz to recorded neurons in order to quantify the physiological parameters listed below. All values were computed at a current injection of rheobase + 20 pA unless otherwise stated.

- Membrane resistance (MΩ): Slope of the line fit to the current-voltage responses for all current steps prior to rheobase current.
- Maximum firing rate (Hz): Mean firing rate during first 200 ms of spiking on the last current step before a reduction in firing rate occurs.
- Maximum action potential rising slope (mV/ms): Greatest value of action potential rising slope between threshold and peak.
- Maximum action potential downslope (mV/ms): Most negative value of action potential falling slope after peak has been reached.
- First action potential latency (ms): Time from initiation of current injection to first action potential peak.
- Action potential threshold (mV): Membrane voltage at which action potential rising slope first crosses 20 V/s.
- After-hyperpolarization (AHP) amplitude (mV): Largest hyperpolarized deflection within 15 ms of action potential. Amplitude was computed as the difference in voltage between action potential threshold and maximal hyperpolarization peak.
- AHP latency (ms): Difference in time between action potential threshold and hyperpolarization peak.
- AHP rise (ms): Time constant computed by fitting a single exponential rise function to the post-AHP repolarization.
- Action potential amplitude (mV): Difference between action potential peak and threshold.
- Action potential halfwidth (ms): Time difference between the rising and falling slope of the action potential, measured at half-amplitude.
- Inter-spike interval (ISI) coefficient of variance: *ISI_stand.dev_*. ÷ *ISI_mean_*.
- First ISI (ms): First ISI.
- Second ISI (ms): Second ISI.
- Firing bias: (*N_spikes_*_,2*nd* 500 *ms*_ − *N_spikes_*_,1*st* 500 *ms*_) ÷ *N_spikes,total_*. Positive values occur for neurons with biases towards late firing; negative values occur for neurons with biases towards early firing.
- ΔAHP (mV): *AHP amplitude*(*last*) − *AHP amplitude*(1*st*) computed at the current step that elicited maximum firing.
- Post-injection voltage step (mV): *Vm_150 ms,post_*_−*injection*_ − *Vm_150 ms,pre_*_−*injection*_ computed at the current step that elicited maximum firing.
- Firing rate accommodation: *ISI*_1*st*_ ÷ *ISI_mean of last 3 ISI_* computed at the current step that elicited maximum firing. <1 indicates ISI slowing; >1 indicates ISI speeding up.
- Amplitude accommodation: *Action Potential Amplitude_mean of last 3_* ÷ *Action Potential Amplitude*_1*st*_ computed at the current step that elicited maximum firing. <1 indicates amplitude shrinking; >1 indicates amplitude growing.
- Halfwidth broadening: *Halfwidth*_2*nd*_ ÷ *Halfwidth*_1*st*_ computed at the current step that elicited maximum firing. <1 indicates narrowing; >1 indicates broadening.

Voltage step for membrane decay: To measure a neuron’s membrane decay constant, voltage responses to 10 trials of a − 50 pA, 1 second current step were averaged. The membrane decay constant was computed by fitting a single exponential decay function to the mean voltage response.

##### Anatomical position

Following recording of each neuron, an image was taken at 4X and 40X of the position of the neuron in the VMHvl. These images were then compared to a mouse brain atlas (Paxinos, 3 ed.), and the location of each neuron in mm from bregma was recorded. Anterior-posterior position relative to bregma was included as a predictor in linear regression models and as an input feature for cluster-based classification of VMHvl neuronal subtypes.

#### Pharmacology

NBQX and D-APV were purchased from Tocris Biosciences (Bristol, UK), and picrotoxin was purchased from Sigma Aldrich (St. Louis, MO, USA). NBQX stock was made at 20 mM and diluted to a final concentration of 10 μM in ACSF; D-APV stock was made at 200 mM and diluted to a final concentration of 50 μM in ACSF; and, picrotoxin stock was made at 25 mM and diluted to a final concentration of 100 μM in ACSF. All stocks were stored at −20°C.

#### Statistical analysis and data display

All analysis was performed offline using custom MATLAB code. For comparisons between two groups, normality was first assessed using the Shapiro-Wilk test. Normal data were compared using an unpaired t-test whereas non-normal data were compared using the Mann-Whitney U test. To determine if anatomical position predicted synaptic responses, a linear regression model was fit to model oIPSC amplitude or paired-pulse ratio as a function of anterior-posterior position. Hierarchical cluster analysis using Ward’s method was used to classify VMHvl “core neurons” based on their physiological properties as well as their anterior-posterior position. (Full list of input features for clustering: membrane resistance, maximum firing rate, maximum action potential rising slope, maximum action potential downslope, first action potential latency, action potential threshold, AHP amplitude, AHP latency, AHP rise, action potential amplitude, action potential halfwidth, ISI coefficient of variance, first ISI, second ISI, firing bias, ΔAHP, post-injection voltage step, firing rate accommodation, amplitude accommodation, halfwidth broadening, membrane decay constant, anterior-posterior location.) We normalized these features via z-scoring and ran principal components analysis across these normalized features. We took the top *N* principal components that contributed to 90% of the explained variance in the dataset and used these principal components as the input for the hierarchical clustering analysis and tSNE. Thordike’s procedure was used to determine the final number of clusters in the hierarchical clustering, and these clusters were used to define VMHvl core neurons as class I or II. tSNE clustering was computed using a perplexity of 8 and cosine distances and was used for the purposes for data display. Data visualizations were created using MATLAB and Adobe Illustrator. VMH anatomical drawings were created via traces of mouse coronal plates from Paxinos (ed 3). Normal data are presented as mean ± standard error of the mean; non-normal data are presented as median with error bars extending along the interquartile range.

### Fiber photometry and optogenetics surgery

At 10-24 weeks of age male animals were anesthetized (isoflurane at 3-5% for induction and 1-2% for maintenance) and the skulls were leveled in a stereotaxic device before viral injections and insertion of optic fibers. For both fiber photometry and optogenetics surgeries, optic fibers were fixed to the skull with Metabond (Parkell), and the animals were allowed to recover for at least two weeks before starting any behavioral experiments.

For fiber photometry experiments, we used 0.50 NA, 400 µm optic fibers (FP400URT, Thorlabs) attached to ceramic ferrules (CF440-10, Thorlabs). For fiber photometry recordings from VMHvl-projecting MPO^VGAT^ neurons, Vgat-ires-Cre mice were injected with AAV2Retro/pAAV-Ef1a-DIO-GCaMP6f (UNC GTC Vector Core, titer of 3.6×10^12) at left VMHvl (AP −4.6 mm, ML − 0.7 mm, DV −5.8 mm, relative to the anterior sinus (AP axis) and the brain surface (DV axis)). Optic fibers were then inserted above the left MPO (the fiber tip was placed at AP −3.1 mm, ML - 0.5 mm, DV −4.3 mm). For fiber photometry recordings from VMHvl shell^VGAT^ neurons, Vgat-ires-Cre mice were injected with AAV5/pAAV-Syn-DIO-GCaMP6f (Addgene, titer of 2.1×10^13) at left VMHvl shell (AP -4.65 mm, ML - 0.8 mm, DV -5.8 mm relative to the anterior sinus and the brain surface). Optic fibers were then inserted above the injected region (the fiber tip was placed at AP -4.65 mm, ML - 0.8 mm, DV -5.3 mm).

For optogenetic activation experiments, we used 0.22 NA, 200 µm optic fibers (FG200UEA, Thorlabs) attached to ceramic ferrules (CFLC230-10, Thorlabs). For optogenetic activation of MPO^VGAT^-VMHvl pathway, Vgat-ires-Cre mice were injected with AAV5/pAAV-EF1a-DIO-hChR2(H134R)-mCherry (Addgene, titer of 1.4×10^13) at left MPO (AP -3.1 mm, ML -0.5 mm, DV -4.8 mm), and optic fibers were inserted above left VMHvl (the fiber tip was placed at AP - 4.6 mm, ML - 0.7 mm, DV -5.45 mm). For optogenetic activation of VMHvl shell^VGAT^, Vgat-ires-Cre mice were injected with the same virus at left VMHvl shell (AP -4.65 mm, ML -0.8 mm, DV - 5.8 mm), and optic fibers were inserted above left VMHvl (the fiber tip was placed at AP -4.6 mm, ML - 0.7 mm, DV -5.45 mm).

### Fiber photometry recording

During fiber photometry experiments, we recorded both the calcium signals and the videos of animals simultaneously using Blackfly S camera, FLIR with SpinView software and RX8 data acquisition board (Tucker Davis Technologies). The camera was triggered by the TTL pulses coming from RX8 at ∼40 Hz and all the timestamps of the camera-triggering pulses were saved along with the calcium signals. GCaMP6f recordings were made through a Doric Lenses photometry system (LEDs at 465 and 405 nm, fluorescence mini cube FMC5_E1(465-480)_F1(500-540)_E2(555-570)_F2(580-680)_S, and Newport Visible Femtowatt Photoreceiver Module NPM_2151_FOA_FC). The system was driven by and recorded from using custom code written for a real-time processor (RX8, Tucker Davis Technologies). GCaMP was excited by driving a 465 nm LED light (∼400 Hz at an intensity of ∼10 μW, filtered between 465 and 480 nm) delivered to the brain through a fiber optic patch cord (MFP_400/430/1100-0.57_0.6m_FCM-MF2.5) and a rotary joint (FRJ_1×1_PT__1m_FCM_0.15m_FCM) The emission fluorescence passed from the brain through the same patch cords and was filtered (500-520 nm), amplified, detected, and demodulated in real time by the system. Demodulated fluorescence signals were saved at a rate of ∼3 kHz.

In the SOAR task, we recorded 20-40 min for each session to record a minimum of 7 poke trials/session. There were 12 to 18 nosepokes in a typical successful session. If the animal performed more than 20 pokes in a session before 20 min, the session was terminated to avoid expensive injury to the reward animal. The delay periods and interaction periods were uniformly distributed between 0 and 1 seconds and between 0.2 and 3 seconds, respectively.

### Closed-loop optogenetic stimulation

For optogenetic stimulation experiments, we recorded both the timestamps of laser on/off and the videos of animals simultaneously using Blackfly S camera, FLIR with SpinView software and RX8 data acquisition board (Tucker Davis Technologies). The camera was triggered by the TTL pulses coming from RX8 at 40 Hz and all the timestamps of the camera-triggering pulses were saved together with the timestamps of laser on/off. A blue laser (460 nm, 7.0-7.2 mW) was connected to a 200 µm diameter patch cord that was fastened to the mice’s implanted fibers via ceramic sleeves. For all stimulation, we delivered 20 ms pulses given at 20 Hz (40% duty cycle). For the social operant task, we used a closed-loop stimulation program to allow stimulating during either interpoke intervals or during the interaction periods. We implemented four different stimulation schedules in total. Three of them were called “IPI 5”, “IPI 20”, and “IPI full” schedules, because the laser turned on at the beginning of interpoke intervals when the reward animal was removed by the actuator. For “IPI 5” and “IPI 20”, the laser kept on for 5 sec and 20 sec, respectively. For “IPI full”, stimulation continued until the next poke, or until 10 minutes had passed without the animal poking. The “dwell” schedule the laser turned on when the reward animal was delivered to the experimental animal and was terminated when the animal was retracted.

### Analysis of fiber photometry and optogenetic experiments results

All analysis of fiber photometry and optogenetic experiments results were performed using custom Python codes (Python version 3.7.3; packages used were scipy version 1.3.0, numpy version 1.16.4, sklearn version 0.21.2, hmmlearn version 0.2.8, and statsmodels version 0.10.0).

### Preprocessing of calcium signal

Each recording session was separately filtered and z-scored before further analyses. Demodulated calcium signals were first downsampled from ∼3 kHz to 40 Hz (the same sampling rate as the camera frame rate) so that each camera frame would have its corresponding calcium signal value. This was done by taking the median of ∼75 data points of the raw calcium signal around the timepoint of each camera frame. The downsampled signals were then saved and used for the further analysis. To make PETH plots and the encoding models, the downsampled signals were first detrended by subtracting the lowpass-filtered signal at 0.005 Hz to remove the long-term signal decay which was especially observable for the relatively small pathway-specific MPO-VMHvl signals. Then the signal was z-scored by dividing by the standard deviation of the detrended signal within first 100 sec of the resident-intruder sessions or the SOAR task sessions, when there were no social stimuli affecting the signal. Finally, the detrended and z-scored signal was lowpass-filtered at 3 Hz before being used in the encoding models. For PETH plots, we calculated the PETHs the detrended and z-scored signal and then lowpass-filtered the final PETH at 3 Hz. For all the lowpass-filtering, we used a butterpass filter made by scipy.signal.filtfilt and scipy.signal.butter with an order of 3.

### Markerless pose tracking for fiber photometry and optogenetics experiments

We performed tracking using the top-view videos for the resident-intruder sessions while we used the side-view videos for the SOAR task because we needed to track the vertical movement of the reward animal in the SOAR task. Also, because we switched from light-color bedding to dark-color bedding while we performed the experiments, we made different training sets for different cage bedding colors to maintain the quality of tracking. We also made separate training sets for the fiber photometry and optogenetic experiments because the patch cords for these experiments had different colors and diameter.

Each of the training sets included 200 to 400 frames. The following points were tracked for the resident-intruder sessions:

- Resident left ear
- Resident right ear
- Resident left eye
- Resident right eye
- Resident nose
- Resident tail-torso interface
- Resident fiber base
- Intruder left ear
- Intruder right ear
- Intruder nose
- Intruder tail-torso interface

For the SOAR task, we additionally tracked the following points:

- Left nose port (i.e. the social port in the fiber photometry and optogenetics experiments)
- Right nose port (i.e. the null port)
- Intruder jacket loop (the interface between the reward animal (intruder) and the tether connected to the actuator)

The training frames were first selected by k-means clustering of each video session in the training set, but 10 to 20 frames showing two animals were manually added for the videos from the SOAR task because most of the camera frames from the SOAR task videos contain only the experimental animal alone during the inter-poke intervals, which occupied the majority of the frames picked up by the k-means clustering. The training was run for 350,000 iterations.

### Feature definition for the encoding model for fiber photometry data

To define the experimental mouse’s posture with respect to the intruder (in the resident-intruder assays) and the nose ports (in the SOAR task), we calculated behavioral features from the tracking data.

For the resident-intruder assays for the fiber photometry animals, we calculated the features below for each frame to use them as inputs to the encoding model:

- Distance between the experimental animal and the intruder: euclidean distance between the experimental animal’s fiber base and the centroid of the intruder body parts (left ear, right ear, nose, and tail-torso interface)
- Change of distance between two animals: the frame-to-frame differences of the distance between the experimental animal and the intruder as defined above
- Velocity of the experimental animal: the distance for which the experimental animal’s fiber base traveled between a frame and the following frame
- Velocity of the intruder animal: the distance for which the intruder’s centroid (based on left ear, right ear, nose, and tail-torso interface) traveled between a frame and the following frame

For the SOAR task for the fiber photometry animals, we calculated the features below:

- Distance between the experimental animal and the social nose port: euclidean distance between the experimental animal’s fiber base and the center of the left-side nose port
- Vertical orientation of the experimental animal: the difference between the vertical coordinate of the experimental animal’s fiber base and of the experimental animal’s tail-torso interface
- Horizontal orientation of the experimental animal: the difference between the horizontal coordinate of the experimental animal’s fiber base and of the experimental animal’s tail-torso interface
- Velocity of the experimental animal: defined in the same way as for the resident-intruder session
- Distance between the experimental animal and the reward animal: euclidean distance between the experimental animal’s fiber base and the intruder jacket loop

When calculating these features, we first excluded the data points where the likelihood of the pose tracking estimates for each of the relevant body parts and linearly interpolated the gaps afterwards using scypi.interpolate.interp1d (scipy 1.3.0).

### Encoding model

In order to capture the contribution of behavioral events to the calcium signal, we fit separate linear encoding models for SOAR task sessions and a resident-intruder session for each mouse. The objective was to model the z-scored calcium signal during 1) one resident-intruder assay or 2) up to three SOAR task sessions with a submissive male intruder with one set of parameters per animal.

The design matrix for the resident-intruder sessions includes continuous representations of the 4 discrete events: attack onset, attack offset, intruder’s fleeing, male intruder entry. Each of these events were convolved with a set of 21 cubic splines spanning from 2 seconds before to 2 seconds after the discrete events. With 21 splines associated with each of the 4 discrete events, we have a total of 84 splines as regressors. In addition, the design matrix includes 4 continuous variables: distance between the experimental and the intruder animal, frame-to-frame change of that distance, velocity of the experimental animal, and velocity of the intruder animal. These continuous variables were first smoothed by a butterpass lowpass filter at 3 Hz with an order of 3, and the two velocities were logarithmized before being included into the design matrix. The final design matrix had 89 columns: 84 for the discrete events, 4 for the continuous variables, and a constant term.

The design matrix for the SOAR task includes continuous representations of the 6 discrete events: null poke, social poke, reward animal entry (as the time when the actuator started moving down to deliver the reward animal), reward animal exit (when the reward animal was no longer available to attack, which was defined as 2 seconds after the actuator started moving up), attack onset, and attack offset. Each of these events were convolved with the same set of 21 cubic splines which was also used for the model of the resident-intruder sessions. In addition, the design matrix includes 6 continuous variables: distance between the experimental and the intruder animal, distance between the experimental animal and the social nose port, the velocity of the experimental animal, the vertical and horizontal orientation of the experimental animal, and the height of the resident animal’s height hanging from the actuator which was calculated based on the recorded actuator up and down movements. The final design matrix had 133 columns: 126 for the discrete events, 6 for the continuous variables, and last one for the constant term.

The encoding model takes the form of y = βX+ε, where y is the calcium signal for a given mouse with dimension of T, which is the number of video frames for the resident-intruder or SOAR task session. X is the design matrix with dimensions of 89xT. β is the set of weights for each cubic spline or continuous variable or the constant term, with dimension of 1×89. ε is the residual. The weights were learned using statsmodels.api.OLS. For every discrete event, we calculated the associated kernel as the sum of the 21 splines, each weighted by the learned regression coefficients associated with that event and the spline.

### Automatized behavioral classification using random-forest classifier

For the large video dataset for the SOAR task sessions of the optogenetic stimulation experiments, we used a supervised random-forest classifier to identify instances of attack based on the hand annotated data. Each side-view video frame during the SOAR task was classified either as attacking or not attacking (sklearn.ensemble.RandomForestClassifier). The initial dataset consisted of video frames from 14 annotated videos, and the frames were split by 9:1 ratio for training and testing datasets. Video frames of the interpoke intervals were excluded from the initial dataset because attacks cannot happen for those frames.

The feature matrix included following features for each video frames: x and y coordinates of the experimental animal’s head, experimental animal’s tail-torso interface, reward animal’s head, and reward animal’s tail-torso interface; the distance between the experimental animal’s head and the reward animal’s head; the distance between the experimental animal’s head and the reward animal’s tail-torso interface; the distance between the experimental animal’s head and the social nose port; the velocity of the experimental animal. The objective matrix was a binary indicator of if attacking was hand-annotated in that frame. The classifier was trained with a max depth of 2 and 100 estimators. The probability threshold for detecting attacking was set to maximize the sensitivity multiplied by specificity to the power of 4 on the training set.

### Hidden Markov Model

We analyzed the behaviors during the SOAR task sessions of the optogenetic stimulation experiments using a hidden Markov model (HMM). The model was built using the Python package hmmlearn version 0.2.8. The input to the model consisted of two variables: the sequence of a binary variable representing attack (the variable was set to 0 or 1 depending on whether the animal was attacking or not attacking at each timepoint, respectively) and the time course of the experimental animal’s distance to the social nose port, normalized by the min-max normalization method. The input dataset was first made with a sampling rate equal to the frame rate of the video recording (∼40 Hz), and then it was downsampled to ∼4 Hz. The model was first trained on the data from the SOAR task with stimulation performed on the “dwell” regimen because the attack behaviors were manually annotated for these experiments.

Although the default setting of hmmlearn initializes the parameters randomly, we found that the resulting HMM occasionally falls into a local optimum solution in which one of the states occupies most of the states depending on the randomly set initial parameters. Therefore, we manually initialized the emission probabilities so that there are at least one state dominating the attacking data points and another state dominating the data points of large values of the animal’s distance to the nose port. We also manually initialized the transition probabilities between states by setting the diagonal elements to be 0.9 to avoid having the model oscillate between two states.

We tested 2, 3, 4, and 5-state models and used the Bayesian information criteria (BIC) to select which model fit the data best using 10-fold cross-validation. We randomly selected 95% of the original dataset and used it as the training dataset, and then calculated the BIC using the remaining 5% of the original dataset as the test dataset. We iterated this process for 10 times and plotted the averaged BICs, and found a elbow at the 3-state model, suggesting that the performance improved from 2-state model to 3-state model while it didn’t from 3-state model to 4-state model, and therefore we used the 3-state model for the further analyses. The BICs were significantly smaller for the 3-state model than for the 2-state model (t(18) = 4.247, p = 0.000485, n = 10 for each model), but were not significantly different between the 4-state model and 3-state model (t(18) = 0.6355, p = 0.533, n = 10 for each model, both unpaired Student t-test).

Using the 3-state model trained by the dataset from the “dwell” stimulation experiments, we assigned hidden states to each SOAR task session in the optogenetic experiments, and compared the transition probabilities between stim trials and sham trials per stimulation regimens and stimulated regions.

### Histology/tissue processing

Following the completion of our fiber photometry and optogenetic experiments, animals were deeply anesthetized with a ketamine/xylazine cocktail injected intraperitoneally, and transcardially perfused with 4% PFA dissolved in 1x PBS. Brains were extracted from the skull and post-fixed in 4% PFA for 12-36 hours and then they were cryoprotected in 30% sucrose for >24 hours and embedded in optimal cutting temperature mounting medium (Fisher Healthcare Tissue-Plus O. C. T. Compound, Fisher Scientific) for freezing over dry ice. Coronal cryosections of the frozen tissue (50 micrometer slices, Leica) were made and stamped directly onto glass microscope slides.

For fiber photometry animals, we performed immunohistochemistry using anti-GFP antibody to visualize the GCaMP proteins in neurons. Slices were first washed with PBS and then PBS+0.4% Triton (PBT). Then slices were treated with blocking buffer (PBT with 10% donkey serum) for 30 minutes and then underwent incubation by anti-GFP antibody (GFP Polyclonal Antibody, Alexa Fluor™ 488, Invitrogen A21311, Lot #2017366) in PBT with 1% donkey serum at 4℃for 12-24 hours. Then slices were washed by PBT for three times, 10 minutes each, followed by the final wash by PBS for 5 minutes. FInally, the slices were coverslipped with mounting medium (EMS Immuno Mount DAPI and DABSCO, Electron Microscopy Sciences, Cat # 17989-98, Lot 180418). For optogenetics animals, the slices were coverslipped after they were mounted onto the slides using the same mounting medium.

After at least 12 hours of drying, slides were imaged with an automatic slide scanner (Nanozoomer S60, C13210-01, Hamamatsu Photonics), and the fiber locations were manually determined based on the coronal images acquired by the scanner.

For anatomical reconstruction of fibers and tetrodes in animals implanted with 16-tetrode shutleDrives, we used iDISCO brain clearing (https://www.cell.com/fulltext/S0092-8674(14)01297-5) combined with confocal volume imaging and atlas registration. Implanted animals were perfused with phosphate-buffered saline (PBS) followed by 4% paraformaldehyde (PFA), and fixed brains were extracted and soaked in PFA overnight followed by three rinses in PBS. Whole brain tissue was then cleared according to the iDISCO protocol, and volume imaged using a SmartSPIM light sheet microscope (LifeCanvas Technologies) with a 3.6x lens. Finally, data were registered to the Princeton Mouse Brain Atlas (https://brainmaps.princeton.edu/2020/09/princeton-mouse-brain-atlas-links/), and fibers and tetrodes were visualized using Neuroglancer.

### Statistical analyses

Parametric tests, including Student’s *t*-test, paired *t*-test, two-sample *t*-test and one-way and two-way ANOVAs were used if distributions passed Kolmogorov–Smirnov tests for normality. For within-neuron tests of firing rate significance, a non-parametric Wilcoxon signed rank test was used since spike rates were often low and not normally distributed. Repeated tests of significance were corrected with a false discovery rate (FDR) correction or Bonferroni correction. Kolmogorov–Smirnov tests were used to compare distributions of poke latencies. For all statistical tests, significance was measured against an alpha value of 0.05. All error bars show s.e.m. No statistical methods were used to predetermine sample sizes, but our sample sizes are similar to those reported in previous publications. Data here was collected and processed in blocks by methodological subtype and animals were assigned to various experimental groups in blocks after task training. Data collection and analysis were not performed blind to the conditions of the experiments.

### Data availability

All source data is available upon request.

### Code availability

All code is available upon request and will be made available on the lab github.

## Author Contributions

A.F. conceived of and designed this study. T.M. performed all photometry and optogenetics experiments. S.O. performed all extracellular in vivo physiology experiments. E.M.G. performed all slice physiology experiments. A.F., T.M., S.O., and E.M.G. analyzed data. W.F. and I.W. contributed to the slice physiology experiments. P.A. and J.H. contributed to animal training and behavioral analysis. T.M., S.O., E.M.G., and A.F. wrote the paper.

---

**Supplementary Figure 1.**
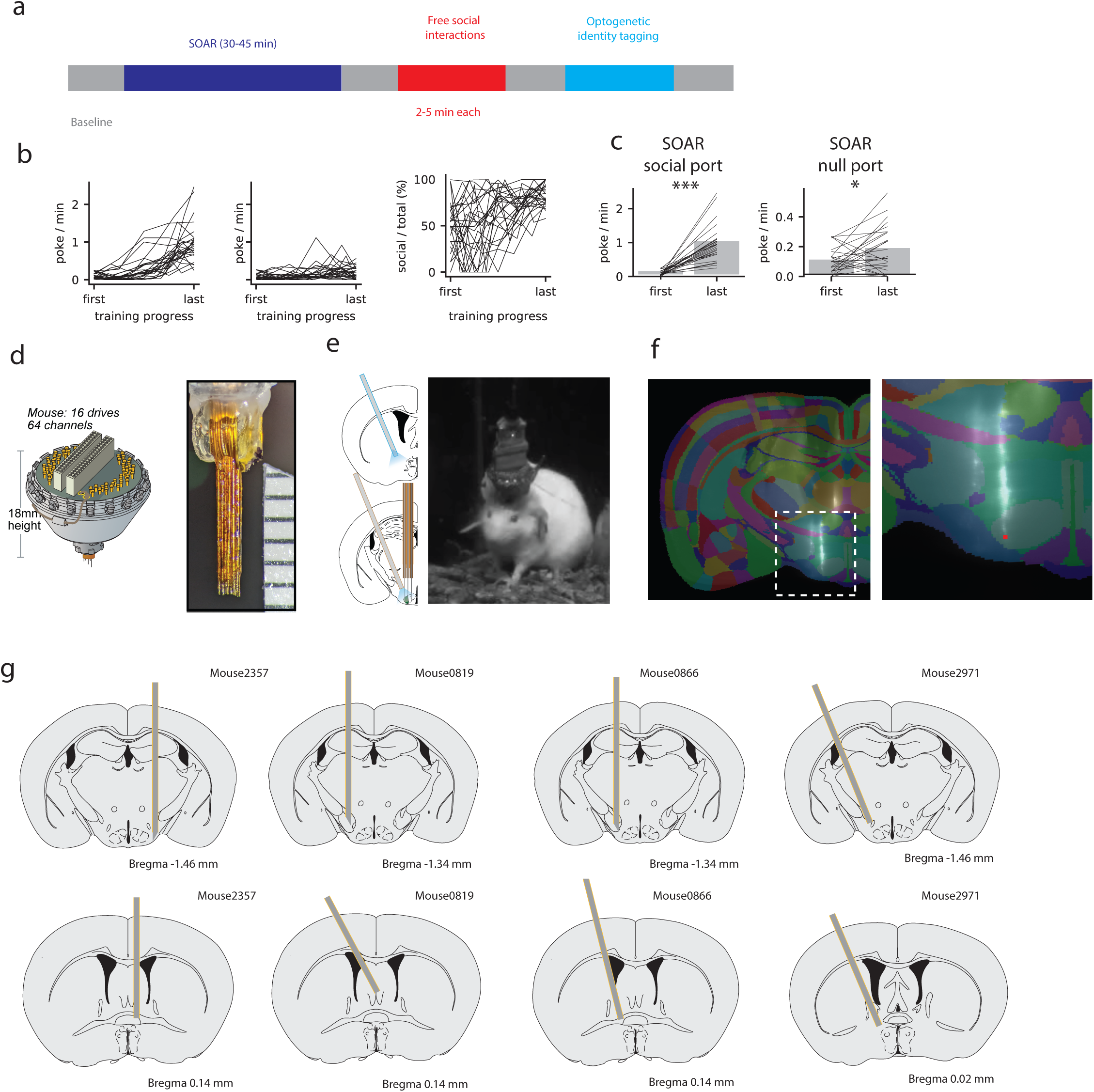
Behavior and histology for chronic physiology during free aggression and SOAR task. **a,** Timeline of example experimental session for chronic recording during behavior. The SOAR task was followed by free social interactions with males, females, and novel objects and identity-tagging experiments were performed last. **b,** Behavior across training of SOAR task. **c,** Mice increased their poke rates to social port and null port across the training. t(36) = −8.184, p = 0.000 for social port; t(36) = −2.390, p = 0.022 for null port; n = 37 mice, paired t-test. **d,e,** Schematic of shuttle drive, implanted fibers and image of implanted mouse **f,** Reconstruction of tetrode locations using iDISCO, example coronal plane of VMHvl (left) and zoomed in reconstruction of a single tetrode track (right). **g,** Fiber locations for implanted animals for identity-tagging experiments.

**Supplementary Figure 2.**
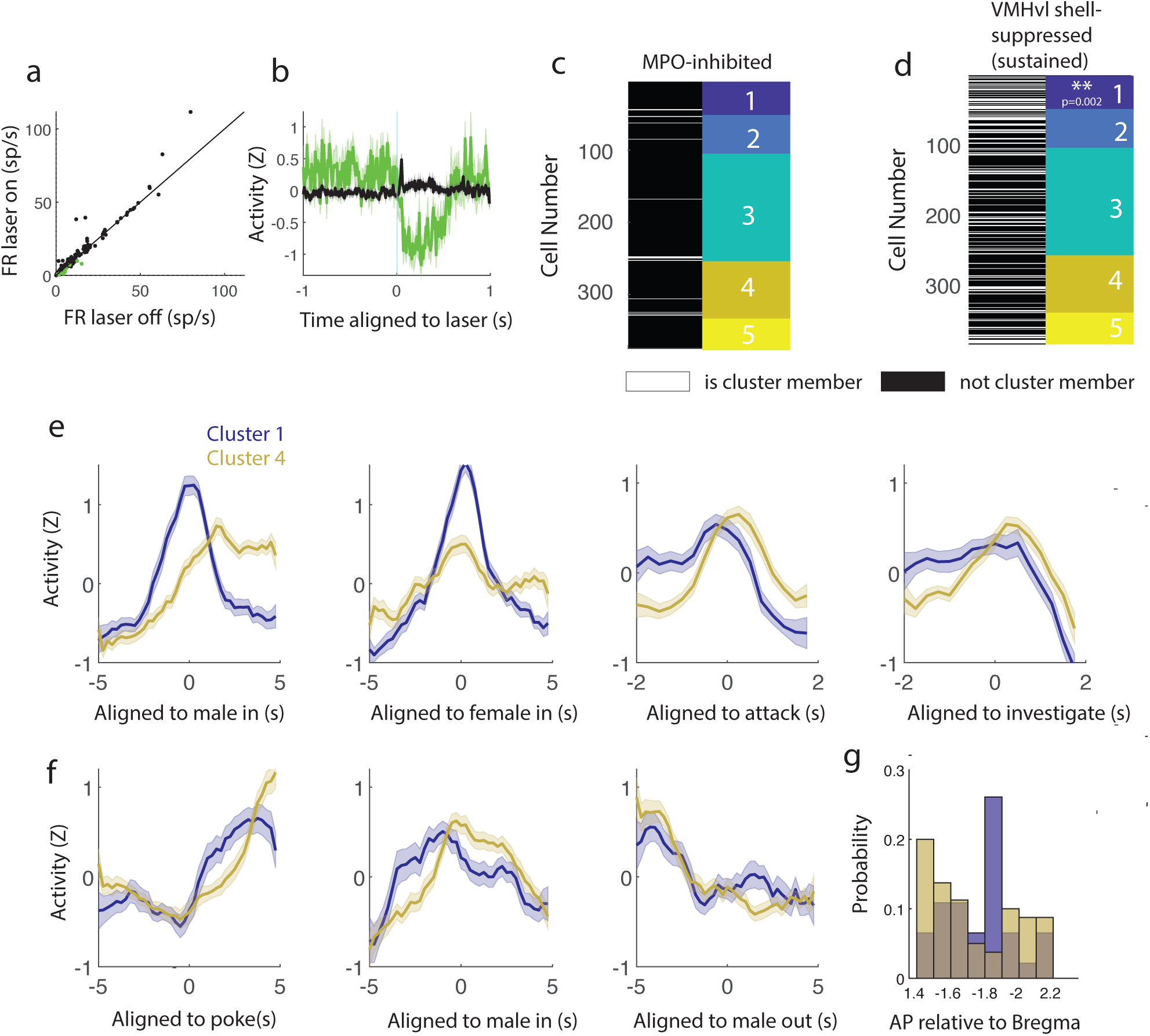
Additional optogenetic Identity tagging experiments. **a-b,** A small number of neurons were identified as being suppressed by inhibition of MPO^vgat+^ (n=16 significant neurons out of 393 neurons, p<0.05 Wilcoxon test with false discovery rate correction). **c,** MPO-suppressed units (n=16 neurons) were not significantly associated with any cluster (p>0.05 for all clusters, Fisher’s exact test with Bonferroni correction). **d,** Cluster identities of VMHvl shell suppressed neurons with less stringent criteria (92/393 neurons) reveals enrichment in cluster 1 (p<0.002, Fisher’s exact test with Bonferroni correction for multiple clusters). **e,f,** Comparison of neural dynamics for cluster 1 (blue, shell-suppressed enriched) with cluster 4 (gold, VMHvl shell enriched) for free behaviors (**e**) and for held out behaviors in the SOAR task (**f**). **g,** Distribution of anatomical positions along the A-P axis of neurons in cluster 1 (blue) and cluster 4 (gold). Neurons in cluster 1 are predominantly in a narrow range in the posterior VMHvl.

**Supplementary Figure 3.**
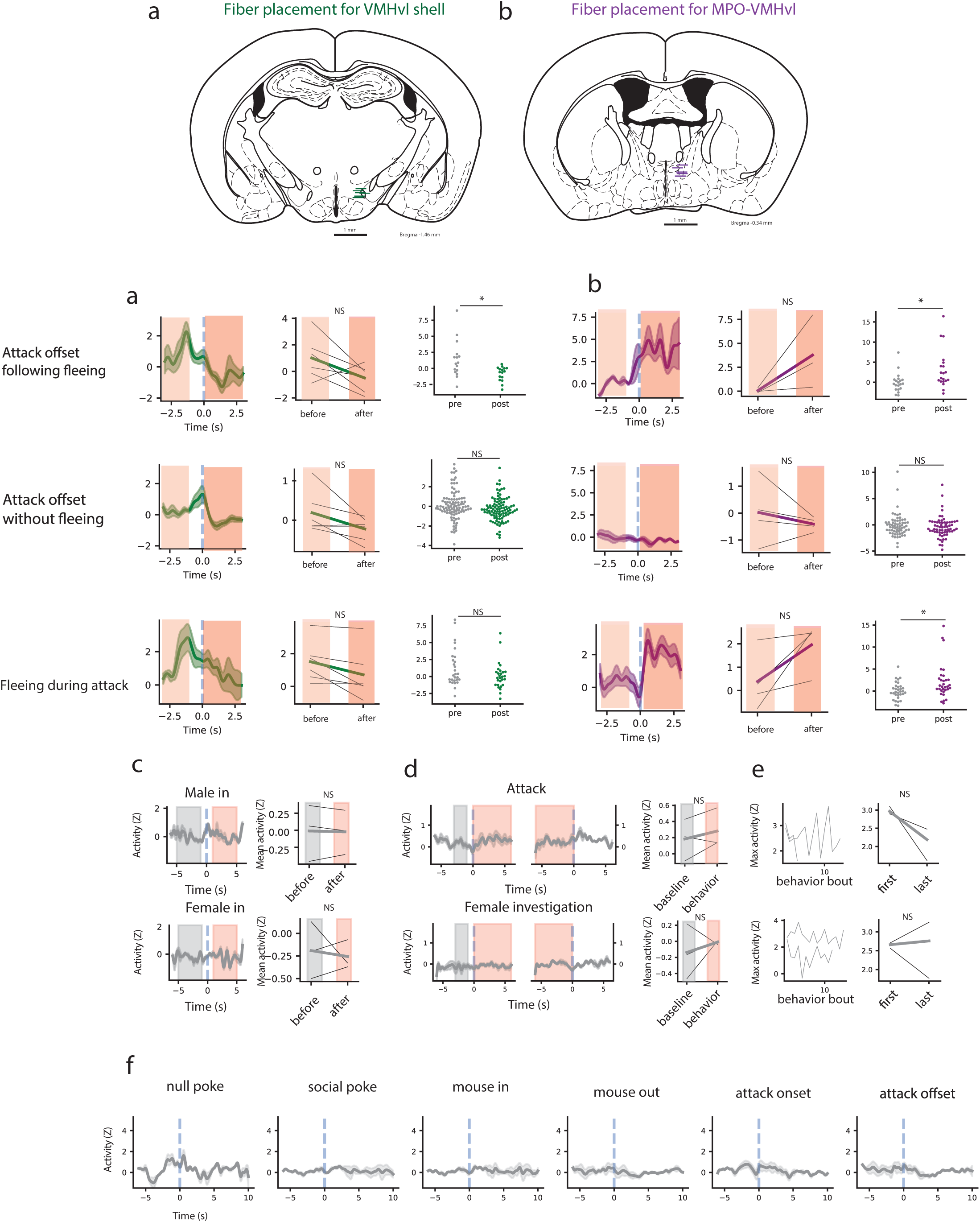
Fiber placement and control experiments for fiber photometry. **a-b,** Locations of fiber tips for the fiber photometry recording for VMHvl shell^vgat+^ (**a**) and MPO-VMHvl^vgat+^ (**b**) scale bars = 1 mm. **c,d,** The GCaMP6f responses to intruder’s fleeing-related events. Top, the GCaMP6f responses to attack offsets happening in <0.5 sec after intruder’s fleeing. Middle, the responses to attack offsets that didn’t happen in <0.5 sec after intruder’s fleeing. Bottom, the responses to intruder’s fleeing during attack. The left column shows the averaged z-scored GCaMP6f activities in the VMHvl shell^vgat+^ and the center column shows the AUCs before versus after the events. The AUC windows are −3 to −1 sec (before) and 0 to 3 sec (after). The right column shows the distributions of AUCs for the same before- and after-window per trial. **e,** Left, average z-scored signal from GFP control animals aligned to the male (top) and female (bottom) intruder entries. Solid lines show mean across mice and shaded regions show standard error. Right, comparisons of AUCs of z-scored signal from GFP control animals before versus after intruder entries. The AUC windows are −5 to −1 sec (before) and 1 to 5 sec (after) around the intruder entries, as shown by the gray(before) and red (after) shadings in the left panels. The thin black lines represent each animal within each group. The AUC after male or female intruder entries was not significantly different from the AUCs before intruder entries for the GFP controls (t(2) = −0.220, p = 0.846 for male entries; t(2) = 0.336, p = 0.769 for female entries; n = 3, paired t-test). **f,** Left, average z-scored signal from GFP control animals aligned to onsets and offsets of attack (top) and female investigation (bottom). Solid lines show mean across mice and shaded regions show standard error. Right, comparisons of AUCs of z-scored signal from GFP control animals for the baseline versus the behavior windows. The AUC windows for the baseline are - 3.5 t o -0.5 sec aligned to behavior onsets (shown by gray shading in the left panels) and the windows for the behaviors are between onsets and offsets of attack (shown by red shadings in the left panels). The thin black lines represent each animal within each group. The AUC during attack against male intruders or during the female investigation was not significantly different compared to the baseline for the GFP control animals (t(2) = −1.095, p = 0.387 for attack; t(2) = - 0.610, p = 0.604 for female investigation; n = 3, paired t-test). **g,** The maximum value of the signal in response to attacks decreases within a resident intruder session. The left column plots the maximum value of the signal during each bout of attack from the first to the last bout within a resident intruder session. The right column compares the maximum values during the first bout versus last bout within a resident intruder session. The thin black lines represent each animal within each group. The maximum value within the first attack against male was not significantly different from the one in the last attack for the GFP control animals (p = 0.183, n = 3, paired t-test). **h,** Averaged signal from GFP control animals aligned to the different events in the SOAR task. Solid lines show mean across mice and shaded regions show standard error. From left to right, each column corresponds to null pokes, social pokes, mouse in, mouse out, attack onset, and attack offset.

**Supplementary Figure 4.**
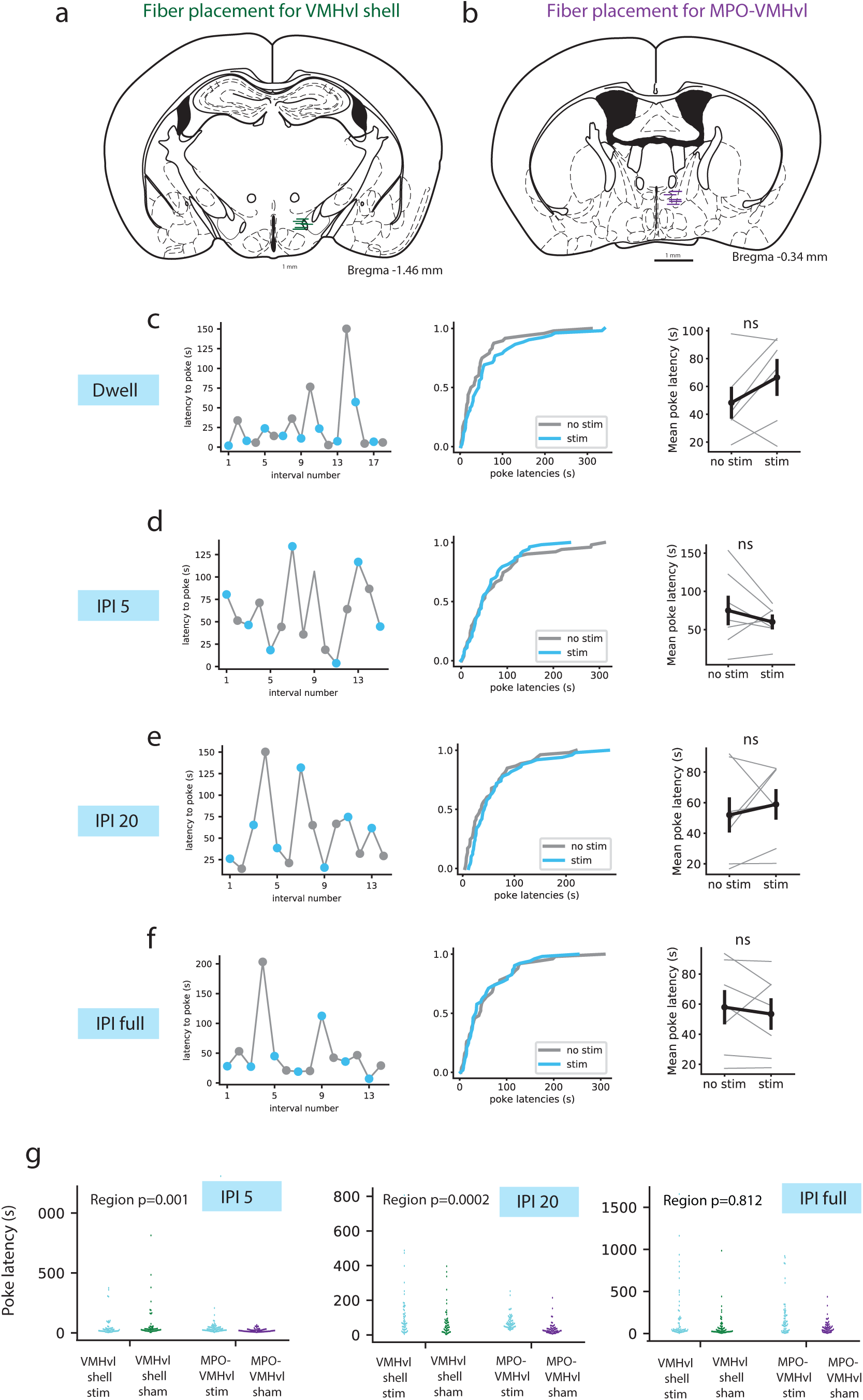
Fiber placement and control experiments for optogenetics. **a-b,** Locations of fiber tips for the optogenetic stimulation for VMHvl shell^vgat+^ (a) and MPO-VMHvl^vgat+^ (b). Scale bars = 1 mm. **c-f,** The effects of optogenetic stimulation during the SOAR task on the poke latencies for the control animals. The left panels show the poke latencies in representative trials for each stimulation schedule. The blue dots represent the poke latencies for stimulation trials and the gray dots represent the poke latencies for sham trials. The center panels show the cumulative fractions of poke latencies across all the control animals. The right panels show the poke latencies averaged across trials per animal. Each thin gray line represents each animal and the solid black line represents the mean across animals with the vertical line showing the standard errors. **c,** The poke latencies were not significantly different between the stimulation trials and the sham trials for the “dwell” regimen for the control animals. per animal comparison: N = 6, t(5) = −1.747, p = 0.141; per trial comparison: N = 53 for stim trials and 49 sham trials, p = 0.390, KS statistics = 0.171, Kolmogorov–Smirnov test. **d,** The poke latencies were not significantly different between the stimulation trials and the sham trials for the “IPI 5” regimen for the control animals. per animal comparison: N = 7, t(6) = 1.069, p = 0.326, paired t-test; per trial comparison: N = 55 for stim trials and 52 sham trials, p = 0.857, KS statistics = 0.109, Kolmogorov–Smirnov test. **e,** The poke latencies were not significantly different between the stimulation trials and the sham trials for the “IPI 20” regimen for the control animals. per animal comparison: N = 7, t(6) = - 0.731, p = 0.492, paired t-test; per trial comparison: N = 51 for stim trials and 54 sham trials, p = 0.130, KS statistics = 0.220, Kolmogorov–Smirnov test. **f,** The poke latencies were not significantly different between the stimulation trials and the sham trials for the “IPI full” regimen for the control animals. per animal comparison: N = 7, t(6) = 0.740, p = 0.487, paired t-test; per trial comparison: N = 53 for stim trials and 51 sham trials, p = 0.827, KS statistics = 0.114, Kolmogorov–Smirnov test. **g,** The poke latencies were significantly different between the VMHvl shell stimulated and MPO-VMHvl stimulated animals for IPI 5 and IPI 20, but not for IPI full regimen. 2-way ANOVA with two factors, the stimulation region and the stim v.s. sham. The region effect was significant for IPI 5 (F_1,1_ = 11.04, p = 0.0010), IPI 20 (F_1,1_ = 14.56, p = 0.00018), but not for IPI full (F_1,1_ = 0.0569, p = 0.81).

**Supplementary Figure 5.**
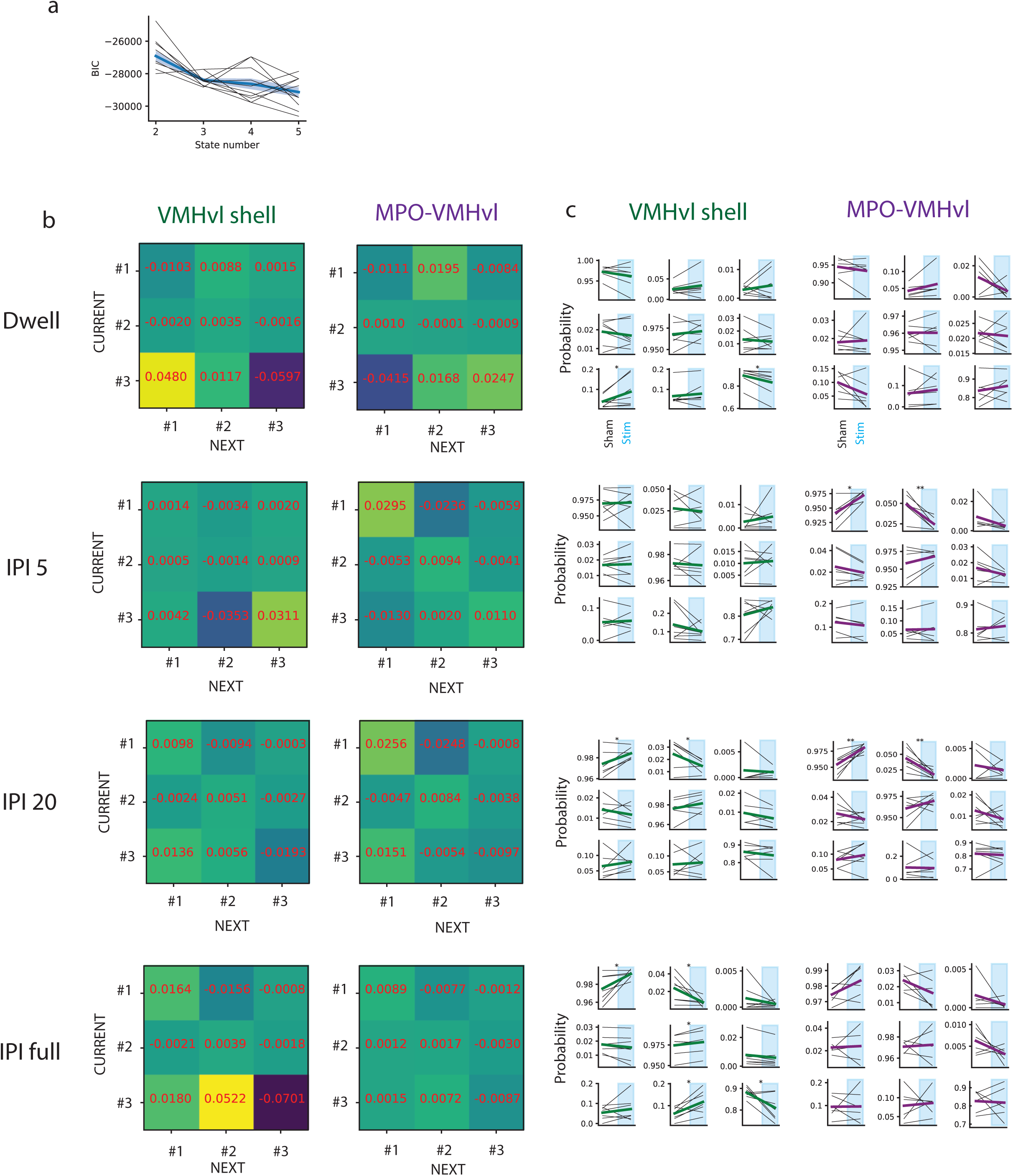
HHM-defined transition probabilities for optogenetic stimulation. **a,** BIC from the models with different numbers of states using 10-fold cross validation. The BICs were significantly smaller for the 3-state model than for the 2-state model (t(18) = 4.247, p = 0.000485, n = 10 for each model), but were not significantly different between the 4-state model and 3-state model (t(18) = 0.6355, p = 0.533, n = 10 for each model, both unpaired Student t-test). **b,** The differences of the state-transition probabilities from the 3-state model between sham and stim trials for the dwell, IPI 5, IPI 20, and IPI full regimens. The rows represent the current states and the columns represent the next states. The positive or negative numbers in each cell of the heatmap indicate that the corresponding transition probability was higher or lower for the stim trials than for the sham trials, respectively. The state #1, #2, #3 are the same as the three states described in Figure 7, and correspond to putative low motivation, high motivation, and attack states, respectively. **c,** The transition probability changes for individual animals between sham and stim trials. There are 9 plots for each pair of stimulation regimen and the stimulation region, and they are organized in the same way as **b**. The black lines represent individual animals and the green or purple lines represent the mean across animals. The t-statistics and the p-values are shown in Table 2.

